# Mtf2 Safeguards Naïve Pluripotency by Restricting Trophectoderm Competence

**DOI:** 10.64898/2026.06.13.727340

**Authors:** Seong-Min Kim, Sohyun Lim, Seong-Kyun Lim, Yeon-Joon Chu, Jiwon Bu, Young-Woo Won, Keun-Tae Kim, Hanbyeol Kim, Cantas Alev, Murim Choi, Chul-Hwan Lee, Hyuk-Jin Cha

## Abstract

The first fate decision in mammalian embryos segregates embryonic and extraembryonic lineages, establishing a largely irreversible transition from totipotency to pluripotency. The mechanisms that enforce this unidirectionality remain incompletely understood. Here, we identify the PRC2.1 subunit Mtf2 as a key epigenetic barrier that safeguards lineage fidelity in naïve embryonic stem cells (ESCs). Despite globally elevated H3K27me3 levels in naïve ESCs, loss of Mtf2 derepresses trophectoderm (TE) regulators, enhances cellular plasticity, generating both ICM- and TE-like populations. Mtf2-deficient cells readily acquire expanded potential with extraembryonic competence and form blastoids without trophoblast stem cell supplementation, whereas loss of the PRC2.2 subunit Jarid2 has minimal effects. Integrated CUT&Tag and transcriptomic analyses reveal that Mtf2 directly represses TE-associated genes in naïve ESCs. Thus, reduced Mtf2 occupancy permits their activation in expanded potential states. Collectively, our findings establish Mtf2 as a central epigenetic barrier that stabilizes naïve pluripotency and restricts inappropriate acquisition of extraembryonic fate.

## INTRODUCTION

During early embryogenesis, cellular potency becomes progressively restricted as development transitions from totipotency to pluripotency ^1^. Totipotent cells can generate both embryonic and extraembryonic lineages, including the trophectoderm (TE), whereas pluripotent cells are developmentally constrained and lack extraembryonic potential ^2–4^. This transition establishes a largely irreversible segregation between embryonic and extraembryonic fates.

The pluripotent state can be subdivided into naïve and primed states, corresponding to the pre-and post-implantation epiblast, respectively ^5^, and is distinguished by fundamentally different signaling dependencies, metabolic programs, and epigenetic landscapes ^6–8^. Notably, recent studies have shown that naïve pluripotent stem cells derived from human and other primates can differentiate into TE lineages in vitro ^9, 10^. Under defined conditions, these cells give rise to trophoblast stem cell (TSC)-like populations and generate blastocyst-like structures (blastoids) from a single cell population without the need for exogenous TSC supplementation, indicating a latent TE differentiation potential within the naïve state ^11–13^.

Beyond conversion from primed to naïve pluripotency ^14, 15^, expanded/extended potential stem cells (EPSCs) ^16, 17^ and a variety of other stem cell states with totipotent-like features have been reported, including 8-cell embryo–like cells ^18^, totipotent blastomere-like cells (TBLCs) ^19, 20^, totipotent potential stem cells (TPSCs) ^21^, totipotent stem cells ^22^, and induced embryo founder cells (iEFCs) ^23^. These systems exhibit varying degrees of both embryonic and extraembryonic competence under defined culture conditions, highlighting that the restriction of extraembryonic fate in pluripotent states is not absolute but can be experimentally overcome.

Cell fate transitions and lineage specification require dynamic reorganization of the epigenetic landscape to precisely regulate the activation and repression of genes necessary for proper embryonic development ^24, 25^. A key regulator of this process is histone H3 lysine 27 methylation (H3K27me1/2/3), a repressive mark critical for establishing facultative heterochromatin ^26, 27^. H3K27 methylation is catalyzed by Polycomb Repressive Complex 2 (PRC2), which comprises the core subunits EZH1/2, EED, and SUZ12 ^28, 29^. PRC2 exists as two subtypes—PRC2.1 and PRC2.2—distinguished by the accessory proteins MTF2 and JARID2, respectively. Both subtypes are required for PRC2 recruitment to chromatin and subsequent deposition of H3K27 methylation in ESCs ^30, 31^, and they carry out distinct, non-redundant functions during embryonic development ^32, 33^. Consistently, H3K27me3 enrichment at pericentromeric heterochromatin increases from the 2-cell to 8-cell stage and is maintained at E3.5, but is reduced in epiblast cells by E5.5, corresponding to the primed pluripotent state ^34^. However, how H3K27me3 constrains lineage options as cells transition from totipotent to naïve pluripotent states remains unclear.

Here, we engineered dual-reporter naïve ESC lines lacking either Mtf2 (M2KO) or Jarid2 (J2KO) to dissect PRC2 subtype-specific roles in pluripotency transitions. We found that Mtf2 and its associated H3K27me3, which is highly enriched in naïve ESCs but reduced in primed ESCs, repress trophectodermal genes. Loss of Mtf2, but not Jarid2, promotes trophectoderm differentiation and enables blastoid formation. These findings identify Mtf2 as an epigenetic safeguard that restricts extraembryonic lineage differentiation and stabilizes naïve pluripotency.

## RESULTS

### High H3K27me3 is accompanied by expression of *Mtf2* in naïve pluripotency

To investigate the role of H3K27me3 in pluripotency states, we first compared isogenic naïve and primed ESCs ^35–38^, examining global H3K27me3 levels along with other pluripotency and self-renewal markers. Naïve ESCs maintained in 2i (GSK3β and MEK1 inhibitors) plus leukemia inhibitory factor (LIF) exhibited substantially higher H3K27me3, Stat3 phosphorylation, and Nanog expression (Fig. 1A), consistent with previous studies ^39^. Transcriptomic analysis revealed that genes encoding PRC2 components, including Jarid2 and Mtf2, were significantly downregulated in primed ESCs (Fig. 1B). This was validated at both mRNA (Fig. 1C) and protein levels (Fig. 1D) across two independent ESC lines (J1 and OG2) ^35^, confirming that Mtf2 and Jarid2 are selectively enriched in the naïve state.

**Figure 1.**
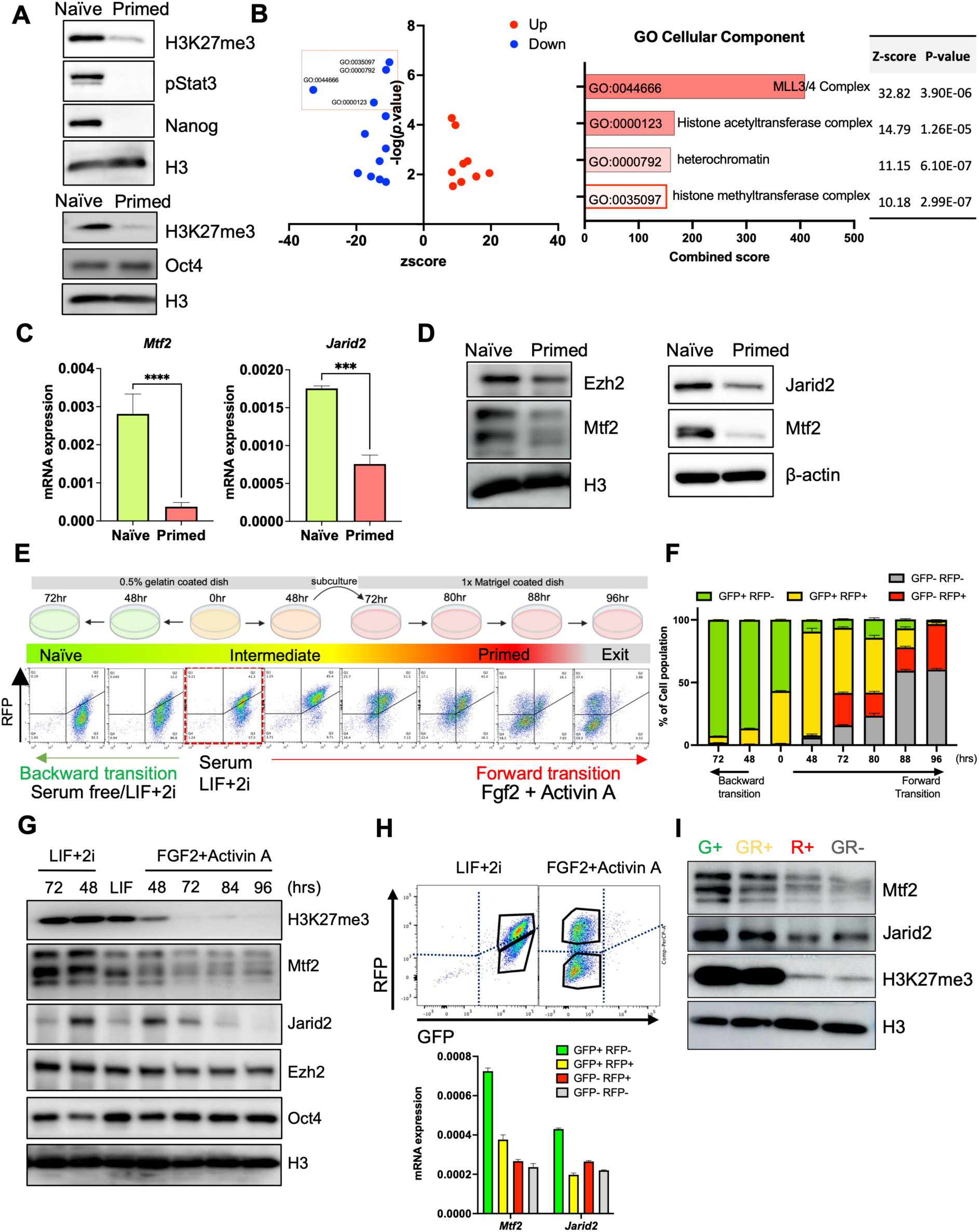
High H3K27me3 is accompanied by expression of *Mtf2* in naïve pluripotency. (A) Immunoblotting analysis for indicative proteins (H3K27me3, pSTAT3, Nanog and OCT4) in naïve and primed mESCs. H3 was used as a loading control. (B) Transcriptome analysis of naïve and primed cells. Volcano plot of Up DEGs and Down DEGs (left), GO analysis of Down DEGs (right). (C) Relative mRNA expressions for two accessory proteins of PRC2, *Mtf2* (PRC2.1) and *Jarid2* (PRC2.2) in naïve and primed ESC cell, t-test, (***, p < 0.001, ****, p < 0.001), (N = 3, n = 3). (D) Immunoblotting analysis for indicative proteins (Ezh2, Mtf2, Jarid2) in naïve and primed mESCs. H3 and β –actin was used as a loading control. (E) Graphical outline for naïve or primed conversion experiments using OG2GOF6 cell line (top), corresponding flow cytometric data (bottom). Naïve conversion was performed under serum-free (N2B27) LIF+2i media, Primed conversion was performed under N2B27 FGF2+ActivinA media (N = 1, n = 3). (F) Quantification of flow cytometry data from left panel (N = 1, n = 3). (G) Immunoblotting analysis for indicative proteins (H3K27me3, Mtf2, Jarid2, Ezh2, OCT4) during conversion toward either naïve or primed pluripotency. H3 was used as a loading control (n = 1). (H) Flow cytometry data of OG2 GOF6 cell line under serum/ LIF+2i condition and under FGF2+Activin A condition. Four distinct populations (GFP+/RFP-, GFP+/RFP+, GFP-/RFP+, GFP-/RFP-) were displayed(up). Relative mRNA expressions for *Mtf2* and *Jarid2* of four indicative populations, (down) (N = 1, n = 3). (I) Immunoblotting analysis for indicative proteins (Mtf2, Jarid2, H3K27me3) of four indicative populations, H3 was used as a loading control (n = 1).

To monitor dynamic transitions between pluripotent states, we employed a dual-reporter ESC system in which GFP and RFP are driven by the distal and proximal Pou5f1 enhancers (Extended data fig.1A), respectively ^40^. Under N2B27 2i/LIF conditions, cells transitioned from a GFP⁺/RFP⁺ intermediate state toward a GFP⁺/RFP⁻ naïve population (hereafter referred to as backward transition), whereas N2B27 FGF2/Activin A treatment promoted progression toward a GFP⁻/RFP⁺ primed state (forward transition) (Fig. 1E). This system enabled tracking of bidirectional state transitions and was validated using established naïve (Extended data fig.1B), primed (Extended data fig.1C), and core pluripotency markers (Extended data fig.1D). Notably, transition to the primed state was accompanied by a marked reduction in global H3K27me3 levels together with decreased expression of Mtf2 and Jarid2 (Fig. 1G). Consistent with this, sorted reporter-defined naïve (GFP⁺/RFP⁻) and primed (GFP⁻/RFP⁺) populations (Fig. 1H), corresponding to the fluorescence profiles (Extended data fig.1E), confirmed that elevated H3K27me3 correlates with robust expression of Mtf2 and Jarid2 in naïve ESCs (Fig. 1I, Extended data figs.1F, and 1G).

### Mtf2 restricts backward transition toward naïve pluripotency

Given that Mtf2 and Jarid2 expression was reduced during forward transition (Fig. 1), we generated CRISPR-mediated knockout ESC lines in the dual-reporter background by targeting exon 11 of *Mtf2* and exon 3 of *Jarid2* (Extended data figs.2A and 2B). Western blotting confirmed successful knockout in two independent clones for each genotype [#2 and #8 for Mtf2 KO(M2KO); #52 and #56 for Jarid2 KO (J2KO)], without major changes in global H3K27me3, H3K9me3, or H3K4me3 levels (Fig. 2A). Flow cytometric analysis of GFP and RFP expression revealed that deletion of *Mtf2*, but not *Jarid2*, significantly increased the GFP⁺/RFP⁻ naïve population under 2i/LIF conditions (Fig. 2B). Consistently, upon 2i withdrawal, WT and J2KO cells rapidly lost GFP expression, whereas M2KO cells retained a robust GFP⁺/RFP- naïve reporter signal (Fig. 2C). These results indicate that Mtf2 loss stabilizes the naïve-like state represented by the GFP⁺/RFP⁻ reporter population.

**Figure 2.**
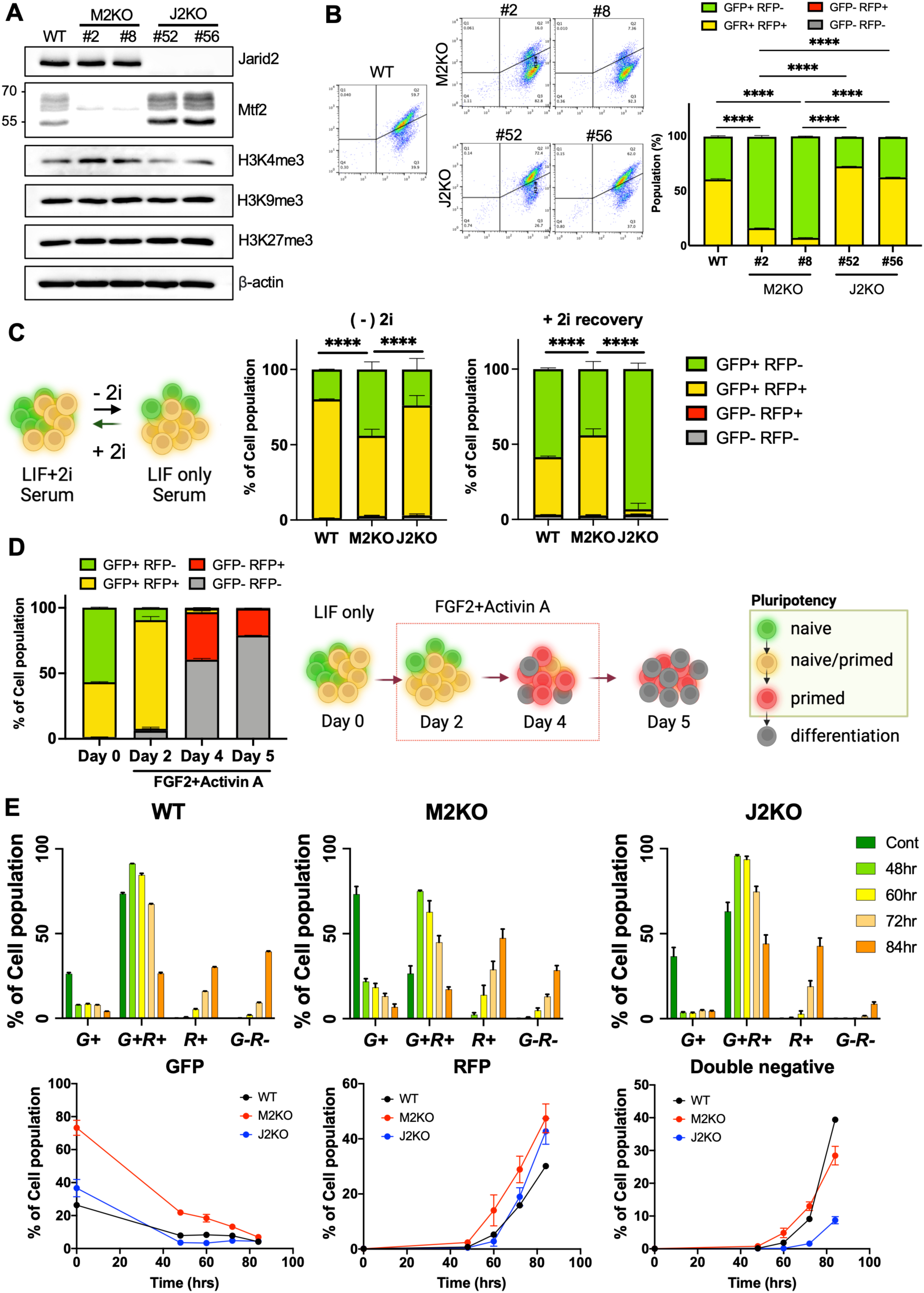
Mtf2 restricts backward transition toward naïve pluripotency. (A) Immunoblotting analysis for indicative proteins (Jarid2, Mtf2, H3K4me3, H3K9me3 and H3K27me3) of WT, M2KO#2, M2KO#8, J2KO#52 and J2KO#56, β-actin was used as a loading control (N = 2). (B) Flow cytometry analysis of WT, M2KO#2, M2KO#8, J2KO#52 and J2KO#56 under serum/LIF+2i culture condition (left). Quantification data of flow cytometry data from left panel (right), Two-way ANOVA, multiple comparisons (****, p < 0.0001), (N = 1, n = 3). (C) Graphical illustration for 2i depletion and re-introduction starting from naïve OG2 GOF6 cells (left). Flow cytometric analysis for 2i depletion [(-) 2i)] and re-introduction (+ 2i recovery) using WT, M2KO and J2KO, (right), Two-way ANOVA, multiple comparisons (****, p < 0.0001), (N = 1, n = 4). (D) Flow cytometric analysis for primed conversion under FGF2+Activin A media using WT (left). Graphical illustration for primed conversion starting from naïve OG2 GOF6 cells (right). (E) Quantification of GFP+/RFP- population, GFP-/RFP+ population, and GFP-/RFP- population of indicative cells at indicative timepoint of primed conversion (top), graph showing gradual changes in GFP, RFP and double negative population of indicative cells(bottom), (n = 3).

We next examined whether loss of Mtf2 or Jarid2 affects forward transition toward primed pluripotency. Upon FGF2/Activin A treatment, WT cells progressed from GFP⁺/RFP⁻ naïve and GFP⁺/RFP⁺ intermediate populations to GFP⁻/RFP⁺ primed cells, followed by accumulation of GFP⁻/RFP⁻ double-negative cells indicative of pluripotency exit (Fig. 2D). M2KO cells underwent forward transition toward the primed state with kinetics comparable to WT and J2KO cells, suggesting that Mtf2 loss does not strongly impair naïve-to-primed conversion (Fig. 2E). In contrast, J2KO cells showed a marked reduction in the GFP⁻/RFP⁻ population, suggesting impaired exit from pluripotency (Fig. 2E). Given that reduction of H3K27me3 during the naïve-to-primed transition facilitates activation of differentiation-associated genes ^41^, these findings suggest that Mtf2 loss preferentially enhances maintenance or re-entry into the naïve state rather than broadly blocking forward transition. Consistent with this model (Extended data fig.2C), J2KO cells exhibited compensatory upregulation of Mtf2 protein (Fig. 2A), which may contribute to reduced pluripotency exit. Together, these data indicate that Mtf2 functions as a barrier to backward transition toward naïve pluripotency, whereas Jarid2 appears to play a distinct role in pluripotency exit.

### Loss of Mtf2 enhances the backward transition toward expanded potential stem cell–like states

Analysis of early embryogenesis transcriptome data (GSE70608) revealed major expression changes during two developmental windows: the oocyte-to-2-cell transition, corresponding to zygotic/embryonic genome activation (ZGA/EGA), and the morula-to-blastocyst transition, corresponding to mid-preimplantation gene activation (MGA) ^24, 42^ (Extended data figs. 3A and 3B). These transcriptional changes were accompanied by dynamic histone methylation remodeling (GSE73952) ^24^ (Extended data fig.3C). Notably, PRC2 components also showed stage-specific expression patterns: *Mtf2* increased at the blastocyst stage, whereas *Jarid2* decreased at the 2- and 4-cell stages relative to oocytes (Extended data fig.3D). Together with the induction of H3K27me3 at bivalent domains in zygotes and morulae ^43^, these observations suggest that H3K27me3 regulation may participate in both genome activation and the transition from totipotency to pluripotency.

Given the limited effects of Mtf2 and Jarid2 depletion on forward transition from naïve to primed pluripotency (Fig. 2), we next examined backward transition toward expanded pluripotent or totipotent-like states. Expanded potential can be induced using a defined protocol that converts naïve ESCs into expanded/extended potential stem cells (EPSCs), which can generate both ESC and trophoblast stem cell (TSC) derivatives—a hallmark of extraembryonic competence ^16, 44^—although they do not fully recapitulate totipotency ^45–47^. Using our dual-reporter system (Figs. 3A and 3F), we observed that the GFP signal associated with naïve pluripotency persisted during EPSC induction but was markedly reduced in intensity (Fig. 3B), resembling the diminished Oct4 distal enhancer activity observed in 8-cell-stage embryos compared with E3.5 blastocysts ^48^. In contrast, the RFP signal associated with primed identity was completely lost (Extended data fig.3E). Established EPSCs also displayed distinct morphology (Fig. 3F) and upregulated totipotency-associated markers, including *Dux, Zscan4, Tcstv1,* and *Tcstv3* (Fig. 3C). Their extraembryonic differentiation capacity was confirmed by the emergence of TSC-like colonies (Extended data fig.3F), induction of TSC markers (Fig. 3G), and further reduction of GFP signal under TSC-inducing conditions (Extended data fig.3H). These results demonstrate that the dual-reporter system captures transition toward an expanded pluripotent state with extraembryonic competence.

**Figure 3.**
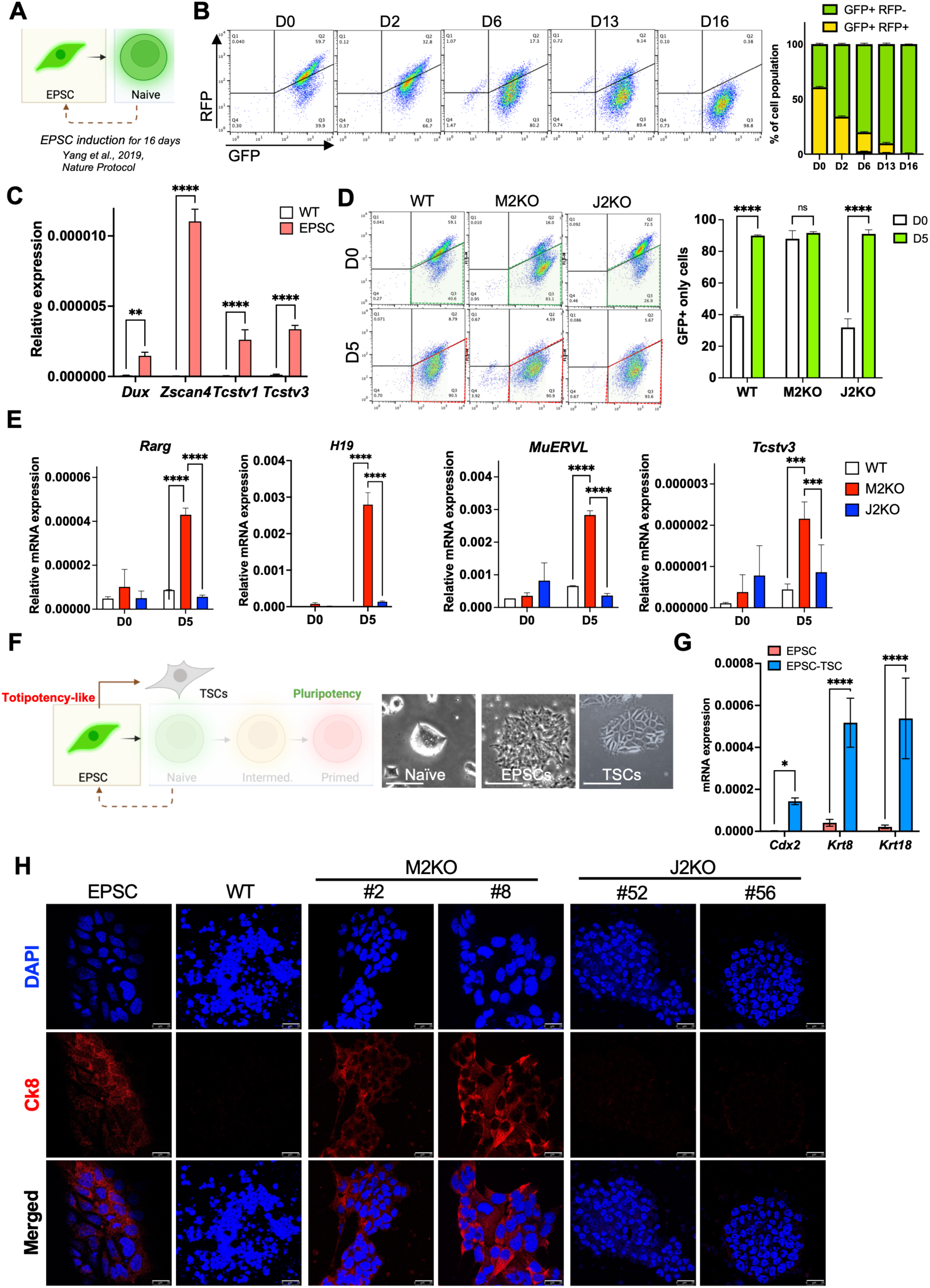
Loss of *Mtf2* enhances the backward transition toward expanded potential stem cell–like states. (A) Graphical illustration of EPSC induction, which is a dedifferentiation from pluripotency to totipotency. (B) Flow cytometry data at indicative days during EPSC induction (left), quantification of GFP+/RFP+ and GFP+/RFP- populations (right). (C) Relative mRNA expressions for totipotency marker genes (*Dux, Zscan4, Tsctv1 and Tsctv3*) of WT and EPSC, multiple t tests, (**, p < 0.01, ****, p < 0.0001), (N = 1, n = 3). (D) Flow cytometry data at indicative days during partial EPSC induction (left) of WT, M2KO and J2KO, quantification of GFP+/RFP- populations (right), multiple t test, (****, p < 0.0001, n.s. for non-significant), (N = 1, n = 3). (E) Relative mRNA expressions of totipotency marker genes (*Rarg, H19, MuERVL, and Tcstv3*) in WT, M2KO and J2KO at day 0 and day 5 of partial EPSC induction, One-way ANOVA, multiple comparisons, (***, p < 0.001, ****, p < 0.0001). (N = 1, n = 3). (F) Graphical illustration for induction toward EPSC and TSCs from Naïve ESCs, and representative brightfield images of Naïve, EPSCs, and TSCs (Scale bars = 100 μm). (G) Relative mRNA expressions for TSC marker genes (*Cdx2, Krt8, Krt18*) of EPSC and EPSC-derived TSC, multiple t test, (*, p < 0.05, ****, p < 0.0001), (N = 1, n = 3). (H) Fluorescent images of 4 days of TSC induction of EPSC, WT, M2KO#2, M2KO#8, J2KO#52, J2KO#56 stained with Ck8 (red) and DAPI for counter nuclear staining.

Unlike naïve mESCs, which are normally restricted from TE differentiation, naïve hESCs can acquire TE potential in vitro ^10^. Previous studies further showed that inhibition of EZH2 enhances TE differentiation from hESCs ^41, 49^. Based on these findings and the high H3K27me3 levels observed in naïve mESCs (Fig. 1), we hypothesized that PRC2-mediated H3K27me3 establishes a refractory state against TE differentiation, with Mtf2 and Jarid2 potentially contributing to repression of TE-associated loci. To test this, we subjected M2KO and J2KO cells, deficient in PRC2.1 and PRC2.2 activity, respectively, to backward transition conditions that partially induce EPSC-like states over five days. Whereas WT and J2KO cells shifted from GFP⁺/RFP⁺ intermediates toward GFP⁺/RFP⁻ naïve cells, M2KO cells retained a stable GFP⁺/RFP⁻ state (Fig. 3D). M2KO cells also strongly upregulated totipotency-associated genes, including *Rarg, H19, MuERV-L,* and *Tcstv3,* compared with WT and J2KO cells (Fig. 3E). Together with the enhanced naïve reversion observed in Fig. 2, these data indicate that M2KO cells are preferentially responsive to backward transitions, including naïve stabilization and EPSC-like conversion, while showing limited effects on forward transition toward the primed state (Fig. 2E).

We next assessed whether this increased backward-transition propensity was accompanied by acquisition of extraembryonic competence. Under TSC-inducing conditions ^50^, which convert GFP⁺ EPSCs into a GFP⁻/RFP⁻ population (Extended data fig.3H), a subset of M2KO cells acquired flattened morphology and showed reduced cell death, whereas WT and J2KO cells formed compact colonies and underwent widespread apoptosis (Extended data fig.3F). The capacity of M2KO cells to generate TSC-like cells was further supported by cytokeratin 8 (Ck8) immunostaining (Fig. 3H) and strong induction of trophectoderm markers, including *Gata2, Gata3, Krt8,* and *Krt18* (Extended data fig.3G). Quantification of the GFP⁻/RFP⁻ population under TSC-inducing conditions further showed a marked increase in M2KO cells (Extended data fig.3I), confirming enhanced acquisition of extraembryonic competence. Collectively, these findings indicate that Mtf2-dependent PRC2.1 activity functions as an epigenetic barrier that suppresses backward transition from naïve pluripotency toward expanded potential and extraembryonic competence.

### Transcriptomic profiling reveals shared extraembryonic competence between M2KO and EPSCs

To further assess the extraembryonic competence of M2KO, we performed single-cell RNA-seq and bulk RNA-seq on WT, M2KO, and EPSCs. Dimension reduction revealed distinct clustering by cell type (Fig. 4A), while principal component analysis (PCA) showed close alignment of M2KO and EPSCs in both single-cell and bulk RNA-seq data (Extended data figs.4A and 4B). Pseudotime analysis showed a pattern highly concordant with principal component (PC) 1 values, revealing a trajectory in which EPSCs and M2KO co-localized and displayed a gradual downregulation of key pluripotency genes including *Klf4, Klf2, Nanog, Esrrb,* and *Zfp42* (Fig. 4B).

**Figure 4.**
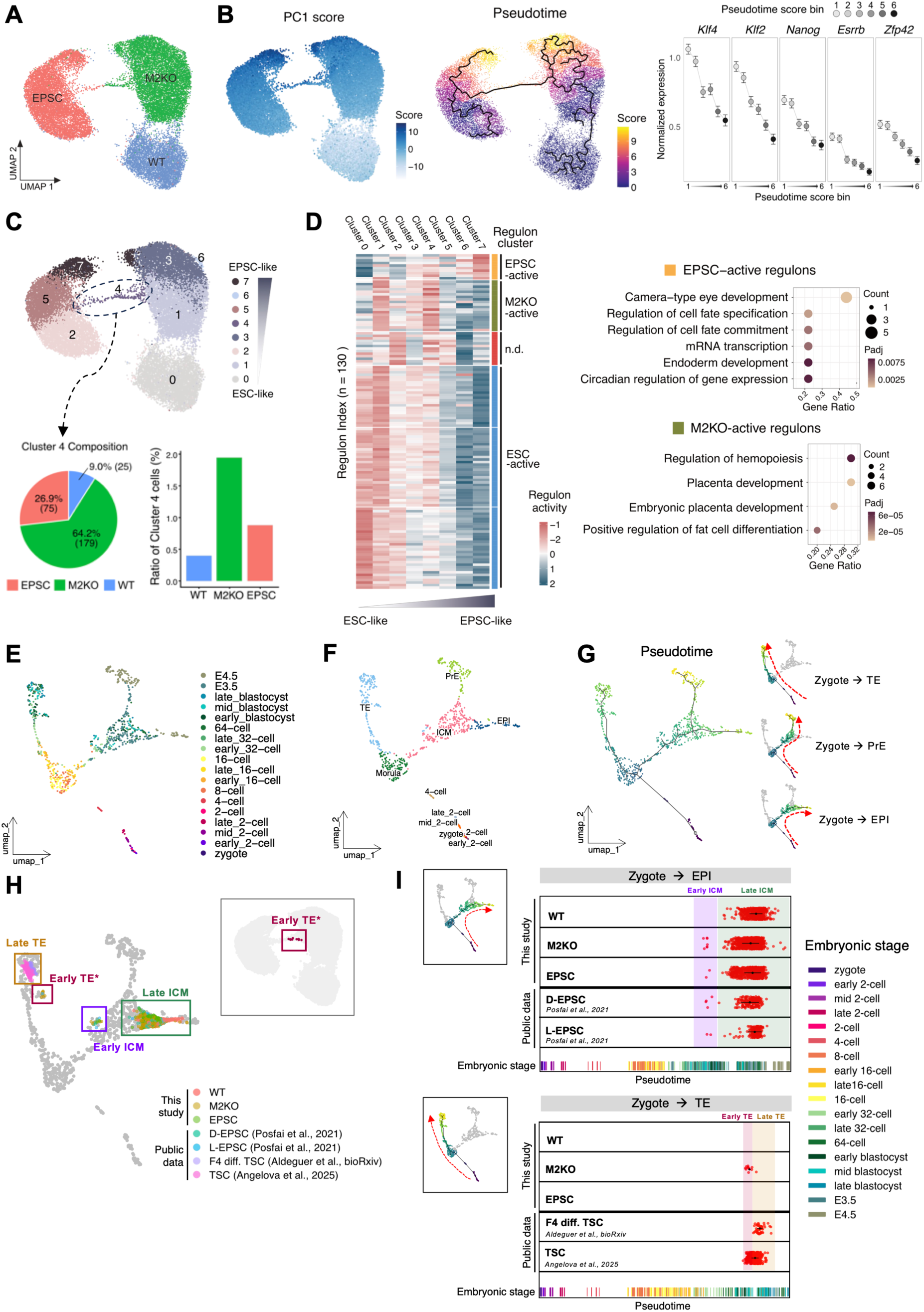
Integrated transcriptomic profiling of M2KO reveals extraembryonic competence. (A) Uniform manifold approximation and projection for dimension reduction (UMAP) of scRNA-seq data, grouped by three different cell types. (B) UMAP plots with cells colored by PC1 values (left) and pseudotime scores (middle). Normalized expression of *Klf4, Klf2, Nanog, Esrrb*, and *Zfp42* in each pseudotime score bin is shown, with error bars representing 99% confidence intervals (right). (C) UMAP showing eight distinct subclusters (top). Quantative distribution of cluster 4 cells across genotypes, illustrated as a pie chart (bottom-left) and a bar plot representing the ratio of cluster 4 cells within each genotype (bottom-right). (D) Relative regulon activities in each cell subcluster shown in (C) (left). Regulons were further grouped (labeled “EPSC-active”, “M2KO-active”, “n.d.”, and “ESC-active”) by k-means clustering (*k* = 6) based on patterns of changes in regulon activities from ESC-like to EPSC-like subclusters. GO terms significantly enriched (*P_adj_* < 0.05) for TFs comprising “EPSC-active” and “M2KO-active” regulon clusters, identified by gene set enrichment test (right). (E) UMAP visualization of integrated *in vivo* single-cell RNA-seq datasets encompassing embryonic stages from zygote to E4.5. (F) Same UMAP as in (E), with cells colored by annotated cell types. (G) UMAP displaying pseudotime trajectories, illustrating sequential lineage segregation into three major developmental branches during early mouse embryogenesis. (H) Projection of our single-cell dataset onto the integrated UMAP shown in (E), alongside publicly available datasets of EPSCs and TSCs. Projected cells were grouped into four distinct clusters; cells assigned to the ‘Early TE’ cluster are highlighted in our UMAP plot (upper right). (I) *in vitro* cell profiles distributed according to their pseudotime along the Zygote-EPI trajectory (top) and the Zygote-TE trajectory (bottom). The segments at the bottom of each plot represent *in vivo* embryonic cells from various developmental stages, arranged by their corresponding pseudotime.

Sub-clustering identified eight distinct cell states (Fig. 4C). To define the biological identity of these sub-clusters, we examined the expression of lineage-specific and developmental potency markers. Notably, cluster 4 showed a localized enrichment of trophectoderm and totipotency-associated genes. Single-cell feature plots confirmed the specific and localized upregulation of key TE markers (*Krt8*, *Krt18*, and *Gata2*) along with the totipotency-related marker *H19* within this bridge-like cluster 4 (Extended data fig.4C), underscoring its extraembryonic transcriptional profile. Having established cluster 4 as a state primed toward the extraembryonic lineage, we next evaluated its cellular composition. Interestingly, composition analysis revealed a marked enrichment of M2KO cells compared to WT (Fig. 4C). Consistent with this, quantification of this cluster 4 proportion within each genotype revealed a specific expansion of cluster 4 in M2KO (1.95%) compared to WT (0.4%) and EPSCs (0.88%) (Fig. 4C). Quantification of transcription factor (TF) activities across these clusters revealed four “regulon clusters” (Fig. 4D). Enrichment analysis revealed that TFs in EPSC-active regulons (with high activity in clusters 6 and 7) were associated with cell fate specification, whereas M2KO-active regulons (with high activity in clusters 1, 3, and 4) were linked to placental development; in contrast, ESC-active regulons (with high activity in clusters 0–2) were enriched for TFs governing pluripotency maintenance and stem cell identity (Fig. 4D and Extended data fig.4D).

We next integrated our single-cell RNA-seq data with published datasets spanning zygote to E4.5 ^51–54^ to position these populations within embryonic development (Fig. 4E). The integrated UMAP recapitulated lineage bifurcations between TE and inner cell mass (ICM), followed by segregation of primitive endoderm (PrE) and epiblast (EPI) (Fig. 4F and Extended data fig.4E). Trajectory reconstruction identified three branches (Zygote–TE, Zygote–PrE, and Zygote–EPI) (Fig. 4G), and projection of additional *in vivo* ^45, 55^ and *in vitro* ^45, 55–57^ datasets confirmed proper alignment (Extended data fig.4F).

We then projected our WT, M2KO, and EPSCs, alongside public EPSC ^45^ and TSC datasets ^56, 57^, which revealed four distinct clusters: ‘Early ICM’, ‘Late ICM’, ‘Early TE’, and ‘Late TE’ (Fig. 4H). Correlation analysis with *in vivo* embryo data indicated: (1) the ‘Late ICM’ cluster—comprising most cells across all groups—resembled EPI; (2) the ‘Early ICM’ cluster—present in M2KO and EPSCs but not WT—showed moderate similarity to EPI and reduced distance to TE, suggesting a less mature ICM state; and (3) the ‘Early TE’ cluster—exclusively observed in M2KO and corresponding to the previously defined cluster 4—aligned with TE populations, though less mature than the ‘Late TE’ cluster, which consisted of bona fide TSCs (Fig. 4H, Extended data figs. 4G and 4H). In line with previous findings, M2KO clearly displayed a population that lies on the ‘Early ICM’ cluster (marked by a box), where our EPSC data and public D-EPSC and L-EPSC data also showed a distinct population (Fig. 4H and Extended data fig. 4H). Furthermore, M2KO also has a population projected on the ‘Early TE’ cluster (marked by a box) (Extended data fig.4H), which further supports the efficient TSC induction of M2KO cells (Fig. 3).

Cluster distributions (Extended data fig.4I) and differential expression analyses—including heatmaps (Extended data fig.4J), representative marker genes (Extended data fig.4K), and gene ontology enrichment (Extended data fig.4L)—further validated that the ‘Early TE’ cluster exhibited a TE-like phenotype, consistent with the trophectoderm-prone differentiation of M2KO cells (Fig. 3). Pseudotime analysis likewise placed M2KO and EPSCs along both the Zygote–EPI and Zygote–TE trajectories, with M2KO cells preferentially enriched toward the TE branch (Fig. 4I). Collectively, these findings demonstrate that M2KO cells acquire extraembryonic competence, manifesting as transcriptional states corresponding to early ICM and incipient TE.

### Mtf2 loss confers self-blastoid formation competence

To functionally validate the acquisition of extraembryonic competence upon Mtf2 loss in naïve ESCs, we examined their ability to form blastoids, which recapitulate key structural and lineage features of blastocysts in vitro ^58^. Blastoids contain both ICM- and TE-like compartments and can be generated either by co-aggregating ESCs with TSCs ^13, 58, 59^ or from a single totipotent-like cell population, yielding “self-blastoids” ^21, 60^. Because self-blastoid formation does not require exogenous TSC supplementation, it provides a functional readout of acquired TE differentiation competence (Fig. 5A). Using an established self-blastoid protocol ^60^, we first compared EPSCs and WT ESCs. As expected, EPSCs generated blastoids, whereas WT ESCs formed GFP⁺ spheroids lacking blastoid-like architecture (Fig. 5B). EPSC-derived blastoids contained Gata4⁺ cells in a salt-and-pepper distribution and Foxa2⁺ cells adjacent to the ICM-like compartment, consistent with PrE-like organization described in blastoids ^13^ (Fig. 5C). We then applied the same conditions to M2KO and J2KO cells (Extended data figs.5A and 5B). Based on cavitation and blastoid morphology criteria (Extended data fig.5C), M2KO cells showed a modest but significant increase in cavitated and blastoid-like structures compared with WT and J2KO cells (Fig. 5D). Immunostaining of representative M2KO-derived structures revealed expression of lineage markers associated with TE (Ck8 and Cdx2), ICM (Sox2), and PrE (Gata4 and Foxa2) compartments (Fig. 5E) ^13, 21, 59, 60^. These findings suggest that loss of Mtf2 increases the propensity of naïve ESCs to acquire blastoid-like, TE-associated features, although this occurs with limited efficiency.

**Figure 5.**
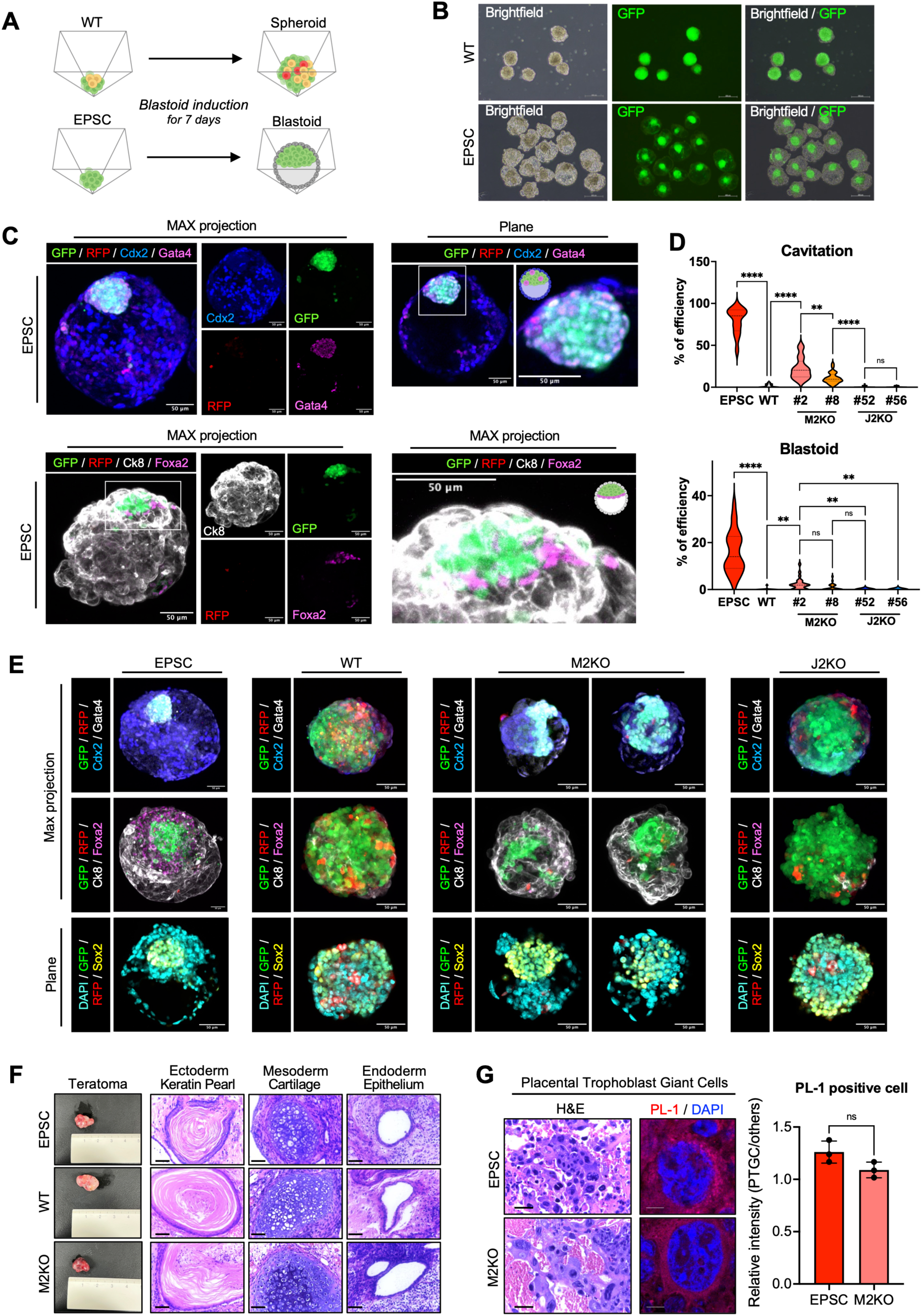
Mtf2 loss confers self-blastoid formation competence. (A) Graphical illustration of the blastoid formation from WT and EPSC cells. (B) Representative brightfield and GFP images of structures on day 7 of blastoid formation from WT and EPSC cells. (C) Immunofluorescence staining of EPSC-derived blastoids on day 7 of blastoid formation for Cdx2 and Gata4 (top), and for Ck8 and Foxa2 (bottom) (scale bars = 50 μm). Composite MAX projection (left), single-channel MAX projection (middle), and composite single plane (top right) or MAX projection zoomed-in images (bottom right). (D) Quantification of ‘Cavitation’ and ‘Blastoid’ structures derived from EPSC, WT, M2KO, and J2KO on blastoid formation day 7. Total 1398 structures of WT, 1372 structures of M2KO#2, 1387 structures of M2KO#8, 1395 structures of J2KO#52, 1399 structures of J2KO#56 were analyzed. Each point represents the percentage of ‘Cavitation’ and ‘Blastoid’ structures in a single imaged field (26 fields in total). RM one-way ANOVA, Multiple comparisons (****, p < 0.0001, **, P < 0.01, n.s. for non-significant), (N = 4, n = 26). (E) Immunofluorescence staining of structures derived from EPSC, WT, M2KO, and J2KO on day 7 of blastoid formation for Cdx2 and Gata4 (top), Ck8 and Foxa2 (middle), Sox2 with DAPI for counterstaining (bottom) (scale bars = 50 μm). (F) Representative images for teratoma mass derived from EPSC, WT and M2KO (left), Representative H&E staining images of teratoma mass derived from EPSC, WT, and M2KO (right). Keratin Pearl for Ectoderm, Cartilage for Mesoderm, and Epithelium for Endoderm (scale bars = 20 μm). (G) H&E staining (left) and Immunofluorescence staining for PL-1 (middle) of Placental trophoblast giant cells from teratoma mass derived from EPSC and M2KO (scale bars = 20 μm, left panels, scale bars = 10 μm, middle panels). Bar graph showing relative PL-1 signal intensity (PTGC / other cells) in EPSC and M2KO-derived teratomas (right). Unpaired t-test (ns, non-significant), (N = 1, n = 3).

To test whether this phenotype was linked to Mtf2-dependent EZH2 activity, we pharmacologically inhibited EZH2 with tazemetostat (iEZH2) ^61^. WT ESCs treated with iEZH2 showed improved blastoid formation by both cavitation and morphology scoring (Extended data figs.5D and 5E). Cdx2 and E-cadherin staining further confirmed TE-like lineage identity and blastoid organization in iEZH2-treated structures (Extended data figs. 5F and 5G).

Finally, we assessed TE differentiation potential in vivo using teratoma formation assays with EPSCs, WT, and M2KO cells (Figs. 5F and 5G). Notably, trophoblast giant cells expressing placental lactogen-1 (PL-1), a marker of trophoblast differentiation, were detected only in teratomas derived from EPSCs and M2KO cells (Fig. 5G), similar to previous observations in teratomas derived from chemically induced totipotent stem cells ^62^. Collectively, these results suggest that Mtf2-dependent EZH2 activity restricts TE differentiation potential in naïve ESCs and acts as an epigenetic barrier to TE competence.

### Integrated transcriptomic and epigenomic analyses reveal Mtf2-dependent regulation of TE competence

To dissect the epigenetic mechanisms underlying the TE-associated phenotypes observed in M2KO, we performed Cleavage Under Target and Tagmentation followed by sequencing (CUT&Tag) for Mtf2, H3K27me3, and H3K4me3, together with bulk RNA-seq in EPSCs, WT, and two independent M2KO clones (M2KO#2 and M2KO#8). Principal component analysis of peak profiles revealed clear segregation by cell type (Extended data fig.6A), demonstrating successful batch correction between the two CUT&Tag experiments. Genomic-feature annotation of called peaks showed that H3K27me3 formed broad domains beyond promoters, whereas H3K4me3 was sharply confined to promoter/TSS regions (Extended data fig.6B). Notably, Mtf2 exhibited a stronger promoter preference than H3K27me3, supporting its role in PRC2 recruitment to promoter-proximal bivalent loci. As expected, strong Mtf2 and H3K27me3 signals were observed at canonical PRC2 targets, such as *Hox* gene clusters, correlating with coinciding with the absence of detectable RNA-seq signal (Extended data fig.6C).

We then focused on “Mtf2-dependent regions”, defined as genomic loci showing a significant loss of Mtf2 CUT&Tag signal in M2KO cells relative to WT (log2 fold change ≤ −1.5). Profiling these loci across WT, EPSCs, and M2KO revealed that H3K27me3 closely tracked Mtf2 occupancy, with a marked reduction in M2KO (Fig. 6A). Notably, Mtf2 signal was also diminished in EPSCs compared with WT, whereas H3K27me3 at the same loci was modestly retained or slightly increased. EPSCs display pronounced bivalency at these sites, with co-occupancy of H3K27me3 and H3K4me3 (Fig. 6A). Consistent with the notion that reduced Mtf2 occupancy at bivalent promoters facilitates gene activation, the overall expression of this gene set was broadly elevated in EPSCs over WT and further upregulated in M2KO (Fig. 6B). Building on our earlier finding that M2KO cells transcriptionally align with EPSCs rather than with naïve WT ESCs (Extended data fig.4B), we next asked whether this resemblance extends to specific gene programs. Quantitative overlap analysis confirmed a strong directional concordance: genes upregulated in M2KO and in EPSCs (each vs WT) overlapped to a far greater extent than expected by chance (representation factor (RF) = 2.8, *P* < 1.7×10^-199^, hypergeometric test; Extended data fig.6D).

**Figure 6.**
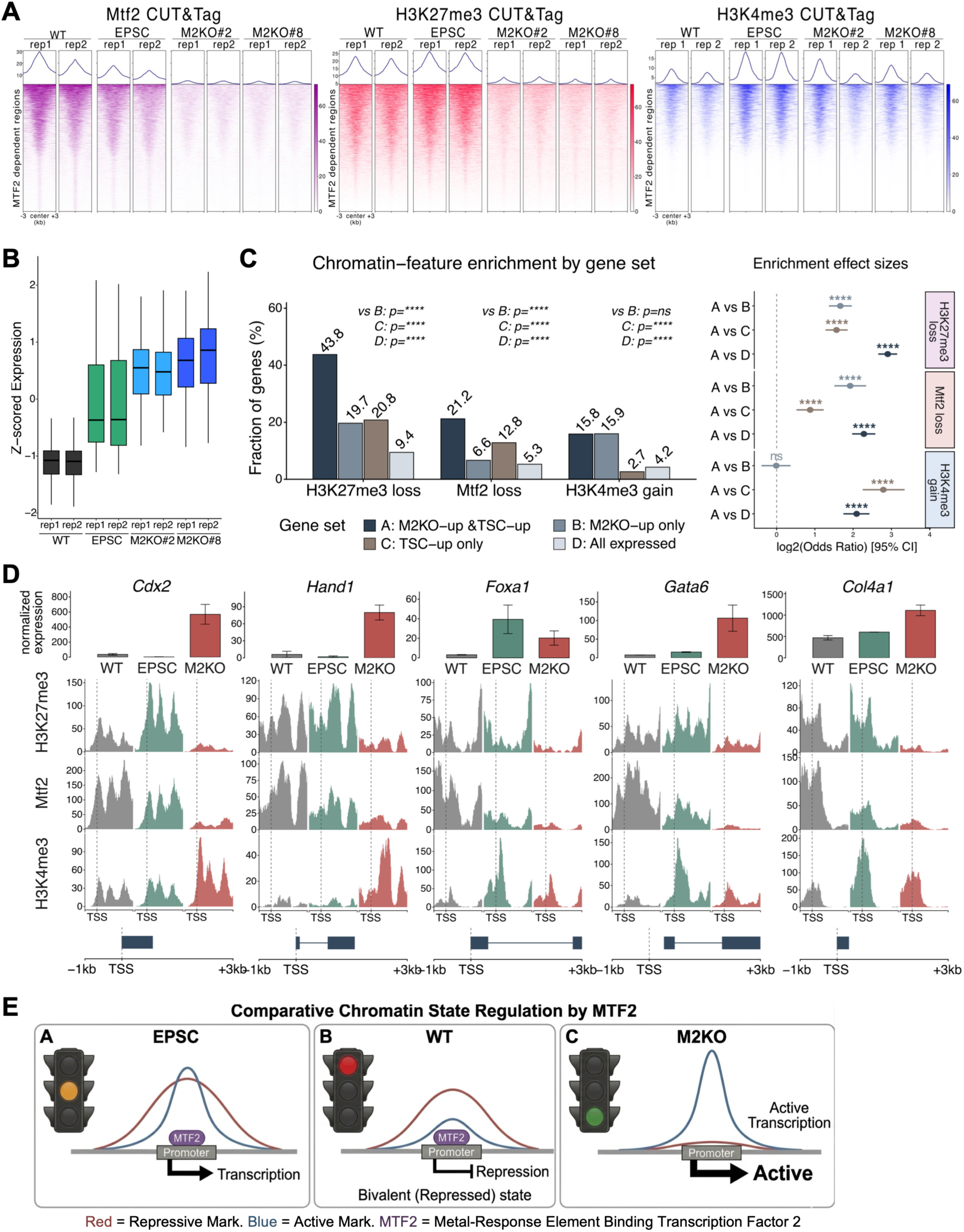
Mtf2 acts as an epigenetic barrier to extraembryonic cell fate. (A) CUT&Tag heatmaps of Mtf2, H3K27me3 and H3K4me3 at ‘Mtf2-dependent regions’ in EPSCs, WT ESCs, and two independent Mtf2-knockout ESC clones (M2KO#2 and M2KO#8). ‘Mtf2-dependent regions’ are defined as Mtf2 peaks with significantly reduced signal in M2KO relative to WT. (B) Box plots showing Z-scored bulk RNA-seq expression of genes associated with Mtf2-dependent regions. Normalized expression values were Z-scaled for comparison. (C) Statistical enrichment of three chromatin features (H3K27me3 loss, Mtf2 loss, H3K4me3 gain) across four gene sets defined within the expressed-gene universe (n=18,198): Set A, M2KO-up DEGs that are also TSC-up DEGs (n=612); Set B, M2KO-up DEGs that are not TSC-up (n=1,568); Set C, TSC-up DEGs that are not M2KO-up (n=2,003); Set D, all expressed genes (n=18,198). Fraction of genes carrying each feature in each set; numbers above bars indicate the percentage (left). Log2 odds ratios (95% CI) from Fisher’s exact tests for Set A vs. Set B/C/D, with row strips colored by chromatin mark (right; **** adjusted *P* < 1×10^-10^, ns, not significant). (D) Representative RPGC-normalized CUT&Tag and RNA-seq tracks for the TSC-related loci *Cdx2*, *Hand1*, *Foxa1*, *Gata6*, and *Col4a1*, shown for the window TSS −1kb to +3kb. Bar plots of normalized expression in WT, EPSC, and M2KO (top). Averaged H3K27me3, Mtf2, and H3K4me3 signals in each cell type (middle; M2KO = mean of clones #2 and #8 across replicates; y-axis scaled per mark to the 99.5^th^ percentile). Gene model (longest transcript) (bottom). (E) Schematic model illustrating Mtf2 function in regulating promoter-proximal bivalency. In EPSCs, Mtf2 facilitates a relaxed bivalent state that permits low-level lineage gene expression, whereas in WT ESCs, Mtf2-mediated H3K27me3 deposition enforces a repressive bivalent state. In M2KO cells, the bivalency is lost, leading to constitutively active chromatin and ectopic lineage gene expression.

To link this transcriptomic output with chromatin changes, we intersected our M2KO transcriptomic profile with a TSC-specific signature (2,615 genes upregulated in TSC vs ESC; GSE159181^63^). Heatmap visualization confirmed that this signature is broadly induced in M2KO cells, recapitulating the expression pattern of TSCs in the reference dataset (Extended data fig.6E). This yielded a core set of 612 TSC-signature genes specifically upregulated in M2KO cells (Set A; Fig. 6C). Relative to the expressed-gene background (Set D, n=18,198), Set A showed a remarkable enrichment of H3K27me3 loss (43.8% vs. 9.4%) and Mtf2 loss (21.2% vs 5.3%) (Fig. 6C, left). To determine whether these changes were a generic consequence of transcriptional upregulation or were specific to the TSC-fate axis, we compared Set A with M2KO-upregulated genes that are not part of the TSC signature (Set B, n=1,568). The enrichment of H3K27me3 loss (OR=3.17, adj.*P* < 1×10^-10^) and Mtf2 loss (OR=3.79, adj.*P* < 1×10^-10^) remained highly significant in Set A over Set B (Fig. 6C, right). In contrast, H3K4me3 gain showed no significant difference between Set A and Set B (15.8% vs 15.9%, OR=0.99, adj.*P* = 0.545; Fig. 6C), indicating that H3K27me3 loss and Mtf2 loss are TSC-fate-specific upstream events, whereas H3K4me3 gain represents a downstream consequence of transcriptional activation. Joint analysis of the three chromatin features confirmed these enrichment patterns at the level of feature combinations: Set A genes more frequently exhibited H3K27me3-loss-containing combinations than Set B, while H3K4me3-gain-only combinations were modestly more frequent in Set B (Extended data fig.6F).

To assess whether the chromatin redistribution observed in M2KO mirrors that occurring during EPSC fate establishment, we applied the same definitions of H3K27me3 loss, Mtf2 loss, and H3K4me3 gain to EPSC-vs-WT contrasts and tested gene-level overlap with the M2KO-vs-WT changes (Extended data fig.6G). Although the absolute numbers of affected genes differed between the two conditions—most strikingly for H3K27me3 loss, where M2KO contained 1,716 affected genes versus 260 in EPSC—the affected gene sets converged at highly significant levels for all three marks. Of the 260 H3K27me3-loss genes in EPSC, 162 (62.3%) were also H3K27me3-loss genes in M2KO (Fisher OR = 17.4, *P* < 1×10⁻¹⁰; Extended data fig.6G). Mtf2 loss and H3K4me3 gain showed similarly significant overlaps (OR = 2.9, *P* < 1 × 10⁻¹⁰ and OR = 3.7, *P* < 1 × 10⁻¹⁰, respectively). These results indicate that M2KO recapitulates—and quantitatively extends—the chromatin redistribution program that accompanies EPSC fate, consistent with a model in which Mtf2 maintains a chromatin barrier whose relaxation underlies EPSC competence. Genomic track analysis of key TSC-related loci – *Cdx2, Hand1, Foxa1, Gata6, and Col4a1* – provided locus-level confirmation of this mechanism (Fig. 6D). These genes, which are bivalently marked or stably repressed in WT ESCs, showed an unequivocal loss of Mtf2 and H3K27me3 in M2KO cells, accompanied by a gain of H3K4me3 at promoters and strongly elevated mRNA expression. In EPSCs, the same loci displayed an intermediate state, with partially reduced Mtf2 and emerging H3K4me3 promoter peaks, in line with their relaxed-bivalency profile (Fig. 6D).

Taken together, these findings establish Mtf2 as a pivotal epigenetic barrier that safeguards naïve pluripotency by recruiting PRC2 to enforce the stable repression of extraembryonic-lineage genes through H3K27me3 deposition. While many of these genes are bivalent in the totipotent morula, they become effectively silenced in naïve ESCs upon stable Mtf2 occupancy. Notably, this barrier is not static but varies across developmental potencies: in EPSCs, partial relaxation of Mtf2 occupancy permits the gradual activation of extraembryonic regulators while preserving bivalency, whereas the complete loss of Mtf2 in M2KO cells leads to widespread derepression and acquisition of extraembryonic competence (Fig. 6E).

### PKC inhibition attenuates Mtf2–PRC2 activity in naïve pluripotency

Because Gö6983-containing naïve culture conditions have been associated with trophectoderm differentiation competence in human naïve pluripotent stem cells ^9, 10^, we asked whether PKC inhibition affects the Mtf2–PRC2 axis. Re-analysis of a published mouse ESC dataset ^64^ showed that PKCi-treated ESCs formed a transcriptionally distinct state compared with conventional 2i/LIF ESCs on PCA (Extended data fig.7A). Notably, both *Mtf2* and *Ezh2* were significantly downregulated upon PKC inhibition (Extended data fig.7B). Consistently, when naïve ESCs were newly derived from primed ESCs in the presence of PKCi (Extended data fig.7C), Mtf2–PRC2 activity toward H3K27me3 was markedly reduced (Extended data fig.7D). We further examined transcriptomic data from human naïve ESCs cultured under PKCi-positive and PKCi-negative conditions across two cell lines (H9 and WIBR3) (GSE153212) ^65^ (Extended data fig.7E). Principal component analysis after batch correction across cell lines and platforms separated PKCi-containing naïve conditions from PKCi-negative naïve and primed conditions (Extended data fig.7F). In this dataset, *EZH2* expression was significantly reduced in PKCi-containing naïve hESCs, whereas *EZH1* remained largely unchanged (Extended data fig.7G). Together, these results suggest that PKC inhibition is associated with attenuation of PRC2-dependent H3K27me3 repression in naïve pluripotency, providing a possible mechanistic link between PKCi-containing culture conditions and increased TE differentiation propensity.

Together, the genetic loss of Mtf2 in M2KO ESCs and the PKCi-induced attenuation of *Mtf2* and *Ezh2* expression converge on the same outcome: a weakened PRC2-mediated H3K27me3 barrier that licenses TE-associated competence. This convergence suggests that the Mtf2–PRC2 axis represents a shared regulatory node through which diverse perturbations—genetic, pharmacological, or species-specific—can modulate the threshold for extraembryonic lineage entry in naïve pluripotency.

## DISCUSSION

In this study, we systematically examined the role of Mtf2, a PRC2.1 accessory protein highly expressed in naïve ESCs and implicated in early embryogenesis. Mtf2 transcription was significantly upregulated at the blastocyst stage, exceeding the corresponding rise in Jarid2 (Extended data fig. 3D). Together with H3K27me3 enrichment at bivalent gene clusters between the morula and ICM ^43^, the association of Mtf2 with H3K27me3 in naïve ESCs (Fig. 1) points to a specific role for Mtf2–PRC2.1 in regulating this developmental transition. Depletion of either Mtf2 or Jarid2 had little impact on forward transitions from naïve to primed pluripotency (Fig. 2), consistent with a recent study showing that PRC2 is dispensable for the naïve-to-primed transition ^66^. In contrast, backward transitions—including naïve reversion and acquisition of TE-associated properties—occurred predominantly in M2KO but not J2KO cells (Figs. 3 and 4). Loss of Mtf2, but not Jarid2, enhanced TSC-like differentiation (Fig. 3), promoted incipient TE-like transcriptional states (Fig. 4), generated cavitated blastoid-like structures at limited efficiency, and produced trophoblast giant cell-like populations in teratomas (Fig. 5). These findings indicate that Mtf2-dependent H3K27me3 safeguards lineage fidelity after blastocyst formation by repressing TE-associated genes that can otherwise be reactivated in permissive states. Embryo-referenced scRNA-seq analyses further showed that M2KO cells display increased propensity toward TSC-like states (Fig. 4 and Extended data fig. 4). CUT&Tag profiling integrated with RNA-seq revealed that Mtf2 contributes to silencing key TE regulators in naïve ESCs, including *Cdx2*, *Hand1*, *Foxa1*, *Gata6*, and *Col4a1* (Fig. 6D). During ZGA, MGA, and the blastocyst transition ^24^, H3K27me3 contributes to lineage fidelity by restricting inappropriate activation of lineage programs ^10^.

Because primed ESCs generally exhibit higher DNA methylation than naïve ESCs ^67^, TE-associated genes required for trophoblast conversion may already be more stably repressed in the primed state. Thus, primed ESCs may rely less on H3K27me3-mediated repression of TE-associated genes than naïve ESCs, providing a possible explanation for the reduced H3K27me3 enrichment observed in primed ESCs (Fig. 1). Naïve mESCs do not readily generate TE, mirroring their behavior in vivo, whereas naïve hESCs established under small-molecule conditions such as Gö6983, a PKC inhibitor in PXGL or t2iLGö medium, can acquire TE differentiation capacity *in vitro* ^10^. This TE competence is further enhanced by pharmacological inhibition of EZH2 ^9, 41, 49^. Rather than challenging these observations, our findings suggest a potential epigenetic basis for this culture-dependent plasticity. Gö6983 treatment has been reported to induce global H3K27me3 loss at transcription start sites in mESCs ^64^. Consistently, our re-analysis of the same mouse ESC dataset ^64^, together with in-house PKCi-treated naïve ESCs, revealed reduced Mtf2–PRC2 activity toward H3K27me3 (Extended data fig.7). Analysis of human naïve ESC datasets further showed that PKCi-containing naïve conditions formed a distinct transcriptional state with reduced *EZH2* expression, whereas *EZH1* remained largely unchanged ^65^ (Extended data figs.7E–G). These observations raise the possibility that PKCi-containing culture conditions partially relax PRC2-associated repression, contributing to TE differentiation propensity in naïve pluripotent states.

Notably, Mtf2 loss was insufficient to fully convert naïve ESCs into TE-competent cells, as self-blastoid formation occurred with limited efficiency. This suggests that Mtf2-dependent H3K27me3 represents one component of a broader epigenetic barrier rather than a standalone determinant of TE fate restriction. Additional repressive mechanisms, including H3K9me3- or H2AK119ub1-associated pathways, may cooperate with PRC2.1 to maintain lineage fidelity in naïve pluripotency. Future studies dissecting how these pathways interact with culture-dependent signaling inputs will help define the broader epigenetic framework limiting TE competence during early mammalian development.

In conclusion, our work identifies Mtf2 as a PRC2.1-dependent safeguard that limits inappropriate activation of TE-associated programs in mouse naïve ESCs. Together with findings from primate naïve PSCs ^68^ and human ESCs ^41, 49^, these results highlight PRC2.1-mediated H3K27me3 repression as a conserved but context-dependent barrier to TE competence during early mammalian development.

## MATERIALS AND METHODS

### Cell lines

Naïve mouse ESCs- J1 (male, ATCC, SCRC-1010, RRID:CVCL_6412), OG2 (male, The Jackson Laboratory, Strain #004654), OG2+/−GOF6+/− (male, derived from Choi et al., 2016) were cultured on 0.5% porcine gelatin-coated plates in either DMEM-based naïve mESC culture media consisting of DMEM high glucose (Gibco) with 15% FBS (Gibco), 1% Glutamax (Gibco), 1% MEM-nonessential amino acids (Gibco), 0.1% Gentamycin (Gibco), 0.1mM β-mercaptoethanol (Gibco), 1000 U/ml mouse leukemia inhibitory factor (mLIF) (Millipore, Merck), 1μM PD0325901 (Biogems) and 3μM CHIR99021-at 37 °C and 5% CO2 incubating condition. Primed mouse ESCs (PJ1, POG2) were cultured on Matrigel (Corning #354277)-coated plates in either EpiSC culture media consisting of DMEM/F12 (Gibco) with 20% KnockOut Serum Replacement (Gibco), 1% GlutaMAX (Gibco), 1% MEM-nonessential amino acids (Gibco), 0.1% Gentamycin (Gibco), 10 ng/ml murine bFgf (Peprotech), 20 ng/ml murine Activin A (Peprotech). Cell lines were routinely passaged every 3 days. Naïve mESCs were treated with Accutase (BD Biosciences) for single cell dissociation and replated at a 1:20 ratio. Primed mESCs were passaged using 1 U/ml Dispase (Gibco) as a colony detachment reagent, with detached colony clumps transferred at a 1:15–1:20 ratio onto Matrigel-coated plates. Culture media was changed daily.

### Establishment and sequence validation of Knockout cell lines

For establishment of Mtf2 KO OG2+/-GOF6+/- cell line, we transfected pRGEN-Cas9-CMV-Puro-RFP (Toolgen-TGEN_OP1) plasmids with sgRNA of Mtf2 cloned into pRG2 (Addgene# 104174) plasmid as previously described ^69^, with the following sgRNA sequence (5’-AGAAGAAGAAGCATTTGTTT-3’) adopted from a previous study ^70^. For establishment of *Jarid2* KO OG2+/-GOF6+/- cell line, pRGEN-Cas9-CMV-Puro-RFP (Toolgen-TGEN_OP1) plasmids were co-transfected with two sgRNAs of *Jarid2* cloned into pRG2 (Addgene# 104174) plasmid, with following sgRNA sequences (5’-CACATACAATCCTTTAAAGG-3’ and 5’-AGCAAGTGTGAATCTACCAC-3’) adopted from the same previous study ^70^. Two sgRNAs were used to remove exon 4 of Jarid2 leading to premature stop codon in Exon 5. The cloned gRNA vectors (3 μg) and Cas9 plasmid vectors (1 μg) were co-transfected into 1×10^6^ mESCs using Lipofectamine 3000 reagent (#L3000-001, Invitrogen). After 24 hours, puromycin selection (2 μg/ml) was performed for 24 hours. Single-colony picking was performed to establish at least two single clones from each Knockout pool. Targeted sequence of each single clone was validated by Sanger sequencing after gDNA isolation through Wizard® Genomic DNA Purification Kit (#A1120, Promega). Knockout efficiency was initially determined by T7E1 assay as previously described ^71^.

### EPSC induction from pluripotent stem cells

EPSC induction was performed as previously described ^16^. The OG2+/-GOF6+/- cells were seeded on 0.5% porcine gelatin-coated dish. After attachment onto plates, cells were cultured in EPSCM – DMEM/F12 high glucose no glutamine (Gibco) supplemented with 20% Knockout Serum Replacement (Gibco), 1% Pen Strep Glutamine (Gibco), 1% MEM-nonessential amino acids (Gibco), 0.1mM β-mercaptoethanol (Gibco), 1,000 U/ml mouse leukemia inhibitory factor (mLIF) (Sigma-Aldrich), 1 μM PD0325901 (Biogems), 3 μM CHIR99021 (Biogems), 4 μM JNK inhibitor VIII (Sigma-Aldrich), 10 μM SB203580 (Sigma-Aldrich), 0.3 μM A-419259 trihydrochloride (Sigma-Aldrich), 5 μM XAV939 (Sigma-Aldrich)- for more than 15 days. The cells were passaged every 2∼3 days. The established EPSCs were maintained in EPSCM as cell line.

### Cell Stock Preparation

Naïve mESCs were cryopreserved using a freezing medium composed of naïve mESC culture medium (see above), FBS (Gibco), DMSO (Sigma Life Science) at a 7:2:1 ratio. Primed mESCs were cryopreserved using mFreSR (Stemcell Technologies). EPSCs were cryopreserved using freezing medium composed of EPSCM (see above), FBS (Gibco), DMSO (Sigma Life Science) at an 8:1:1 ratio. Cells were resuspended in the freezing medium at a density of 0.5-1 x 10^6 cells/mL and transferred to cryovials. Cryovials then moved to −80 °C deep freezer using freezing container overnight before moved to liquid nitrogen for long-term storage.

### Cell sorting of OG2+/-GOF6+/- cell line based on GFP and RFP signal

Cells were dissociated into single cells using Accutase (BD Biosciences), followed by PBS wash x2. Cells were resuspended in PBS and sorted using FACS Aria III cell sorter (BD Biosciences). J1 (mouse naïve ESCs) cells were used as double negative sample for compensation. OG2 (mouse naïve ESCs) cells were used as GFP only positive sample for compensation.

### Induction of Trophoblast Stem Cells

Induction of trophoblast stem cells was performed as previously described ^50^. Cells were seeded on a 0.5% porcine gelatin-coated dish and cultured on LIF+2i media. As TSC media is developed for human cells, we adopted partial induction for up to 4 days. After two days of seeding, media was switched to TSC medium – DMEM/F12 supplemented with 0.1mM 2-mercapto ethanol (Gibco), 0.2% FBS (Sigma-Aldrich, ES-009-B), 0.5% Penicillin/Streptomycin (Gibco), 0.3% BSA (Sigma-Aldrich), 1% ITS-X (Gibco, 51500), 1.5 μg/ml L-ascorbic acid (Sigma-Aldrich), 50 ng/ml mEGF (Gibco, PMG-8041), 2 μM CHIR-99021 (Biogems), 0.8 mM Valporic acid (Tocris, 2815), 0.5 μM A83-01 (Tocris), 1 μM SB431542 (Abcam), and 5 μM Y-27632 (Gibco) – for 4 days.

### Blastoid induction

Blastoid induction was performed as previously described ^60^. Cells were dissociated into single cells using Accutase (BD Biosciences). AggreWell 400 (STEMCELL Technologies) was prepared according to the manufacturer’s instructions. The blastoid formation medium consisted of 25% TSC basal medium (as previously described ^60^), 25% of NDiff227 (TAKARA), and 50% EmbryoMax Advanced KSOM (Sigma-Aldrich), 12.5 ng/ml mFgf4 (R&D systems), 0.5 ng/ml Heparin (STEMCELL Technologies), 3 μM CHIR99021 (Biogems), 5 ng/ml BMP4 (R&D systems) and 0.5 μM A83-01 (Tocris). The dissociated cells were resuspended into the blastoid formation medium supplemented with 2 μM Y27632 (Gibco). Total 30 cells/μWell were seeded onto the prepared AggreWell 400. Total media volume was adjusted to 1 ml by adding blastoid formation medium supplemented with 2 μM Y27632 (Gibco) before cell seeding. The cell seeding day was regarded as day 0 of blastoid formation. The plate was then centrifuged at 800g for 3 min and placed at 37 °C and 5% CO_2_ incubation condition. On day 1, the medium was replaced with fresh blastoid formation medium without Y27632. Subsequently, the medium was changed every 2-3 days. For the media changes, half of the total media volume was gently aspirated and replaced to avoid loss of the structures. On day 7 of blastoid induction, the structures were harvested using a 1000 μl pipette. The pipette tips were cut using sterile scissors to create a broader tip. To avoid attachment and loss of the structures, the cut pipette tip and 6-well plates were coated with 1% BSA in PBS. The blastoids were gently harvested and collected into 1.5ml tube coated with 1% BSA in PBS, followed by 1% BSA in PBS wash x2. The structures were fixed in 300 μl of 4% PFA for 15 min at RT. The fixed structures were stored in 1% BSA in PBS at 4°C for up to 1 month.

### Naïve conversion and primed conversion of OG2+/-GOF6+/- cell line

5×10^5^ Cells were seeded onto 0.5% gelatin coated dish in naïve culture media (supplemented with FBS, see above), a day before the naïve conversion or primed conversion. For naïve conversion, cells were cultured in serum-free naïve culture media-Ndiff227 supplemented with 0.1% Gentamycin (Gibco), 1,000 U/ml mouse leukemia inhibitory factor (mLIF) (Millipore, Sigma-Aldrich), 1 μM PD0325901 (Biogems) and 3 μM CHIR99021 (Biogems) for up to 3 days. For primed conversion, cells were cultured in EpiLC media ^72^, Ndiff227 supplemented with 0.1% Gentamycin (Gibco), 20ng/ml ActivinA (Peprotech), 12ng/ml bFgf (Peprotech) up to 96 hours. After 48 hours of primed conversion, the cells were passaged onto Matrigel (Corning# 354277) - coated dish. The cells were analyzed by flow cytometry at indicative timepoints during naïve conversion or primed conversion.

### Public data analysis

Previous single-cell transcriptome sequencing data from somatic cell nuclear transfer embryos transitioning from oocyte to two-cell stage was downloaded from the Gene Expression Omnibus (GEO)^73^ repository (GSE70605) using sratoolkit v3.0.7 (https://github.com/ncbi/sra-tools/wiki/01.-Downloading-SRA-Toolkit). The raw data in FASTQ format was aligned to the mm10 using STAR aligner v2.7.10a^74^. Subsequent sorting and indexing of aligned reads were performed using samtools ^75^ with default parameters. The Subread package ^76^ was employed to generate a count matrix, which was analyzed using the DESeq2 package ^77^. For downstream analysis, only genes with non-zero counts in a minimum of four samples were included. Gene counts underwent variance-stabilizing transformation (vst) normalization using the default parameters prior to principal component analysis (PCA).

Chromatin immunoprecipitation sequencing (ChIP-seq) data for pre-implantation embryos was downloaded from GEO (GSE73952) ^24^ and processed following the same initial steps as the transcriptome data. Mapped reads were subjected to peak calling using MACS2 ^78^ with following parameters (‘-B -g mm -f BAMPE -q 1e-2 --broad --nomodel --SPMR’). To analyze combinatorial histone mark patterns for H3K4me3 and H3K27me3, ChromHMM v1.25 ^79^ was employed according to the provided instruction ^80^. Emission labels identified by the ChromHMM LearnModel function were annotated into four biological categories: “unmarked”, “H3K27me3 only”, “H3K4me3 only” and “bivalent”. Genomic regions annotated as ‘unmarked’ in at least one sample were excluded from further analysis. For PCA, each biological label was further converted into numeric value (i.e., “H3K27me3 only” as +10, “H3K4me3 only” as −10, and “bivalent” as 0). PCA and subsequent visualizations were conducted using the factoextra R package (https://github.com/kassambara/factoextra), specifically the fviz_pca_ind function.

### RNA isolation and quantitative RT-PCR analysis

Total RNA was extracted using Easy-BlueTM total RNA isola:on kit (iNtRON Biotechnology). cDNA was synthesized using 5x PrimeScript TM RT mix (TaKaRa) for reverse transcrip:on. Using 2x TB-Green premix (TaKaRa), quantitative real-time PCR was performed on a LightCycler-480II (Roche). The gene expression data was normalized using *Rn18s* as an internal loading control.

### Immunoblotting analysis

The whole cell lysate was extracted using RIPA buffer (Biosesang) supplemented with 1 μM protease inhibitor and 10 μM sodium orthovanadate, followed by 1 hour incubation on ice and centrifugation. Protein concentration was determined using the Pierce BCA protein assay Kit (#23225, Thermo Fischer Scientific). The protein samples were mixed with 5x SDS-PAGE loading buffer (#SF2088-110-00, Biosesang) and heated at 100°C for 10 minutes. Total protein (10∼20 μg) was separted on either 7.5% or 10% SDS-PAGE gels, then transferred to Methanol-activated PVDF membranes. The membranes were blocked with 5% skim milk in TBS-T for 1hour at room temperature, followed by overnight incubation at 4°C with primary antibodies (1:500 ∼ 1:1000) diluted in TBS-T containing 1% sodium azide. After washing, membranes were incubated with secondary antibodies (1:10000 in TBS-T) for 1 hour at room temperature. The protein bands were visualized by Chemi-Doc using the Miracle-Star (#16028, iNtRON Biotechnology) chemiluminescence detection kit.

### RNA extraction for bulk RNA-seq

For bulk RNA-seq, the old medium was removed and washed with 1x PBS twice before placing the plate on ice. Next, 1 ml of TRIzol (invitrogen) was added to the plate and incubated on ice for 2-3 min to facilitate cell detachment. The TRIzol with cells was transferred to a 1.5ml tube and incubated for 5 min at room temperature. 0.2 ml of chloroform (Sigma) was added to transferred TRIzol and shaken vigorously for 15 sec and incubated for 3 min at room temperature. After incubation, sample was centrifuged at 12,000g for 15 min at 4°C to separate the sample into three phases. The upper aqueous phase (∼0.5-0.6 ml) was carefully transferred to a new 1.5ml tube. And 0.5 ml of isopropanol (Duksan) was added and shaken vigorously or vortexed and incubated for 10 min at room temperature. After incubation, sample was centrifuged at 12,000g for 10 min at 4°C and the RNA pellet was washed with 1 ml of 70% ice-cold ethanol (Sigma). Finally, the RNA pellet was dissolved in 20-30 μl of DEPC-treated water (invitrogen), the RNA concentration was measured, and the sample was stored at −80°C until use.

### Cell and library preparation for RNA-seq analyses

Libraries for bulk RNA-seq were constructed using the TruSeq Stranded mRNA Library Prep Kit. For single-cell RNA-sequencing, cells cultured in 60 mm dishes were washed with pre-warmed Dulbecco’s Phosphate-Buffered Saline (DPBS) (Gibco). Cells were then dissociated using Accutase (BD Biosciences) after a 3-minute incubation at 37°C. Following dissociation, cells were centrifuged and resuspended in DPBS for single-cell library construction. Library construction for single-cell RNA-sequencing was performed according to the 10X Chromium Next GEM Single Cell 3’ Reagent Kits v3.1 instructions. For each sample, 10,000 cells were targeted, and 10 cycles of amplification were performed for cDNA and library preparation. The libraries were sequenced on the Illumina NovaSeq6000 platform.

### Bulk RNA-seq data processing

For bulk RNA-seq data, adaptor trimming was performed using Trimmomatic ^81^, and trimmed reads were aligned to the mm10 reference genome using HISAT2 ^82^. Aligned reads were processed using the subread package ^76^ to generate count matrices. DESeq2 ^77^ was used for further analysis, including normalization and differential expression analysis. Genes covered by at least 10 reads in more than two samples were included. Differentially expressed genes (DEGs) were identified with an adjusted *P* < 0.05 and a |log2FoldChange| > 1. Gene Ontology (GO) analysis for the DEGs was performed using clusterProfiler ^83^ with the enrichGO function. The statistical significance of the overlap between EPSC and M2KO DEGs was determined using a hypergeometric test. The representative factor (RF) was calculated as:

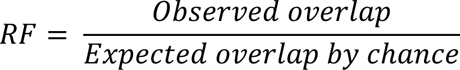

where the expected overlap was derived from the total number of genes detected across all bulk RNA-seq samples (*N* = 18,198). Gene Set Enrichment Analysis (GSEA) was performed using the fgsea ^84^ package v1.28.0 with the MSigDB ^85^ C5 (Biological Process) collection. The ranking metric was defined as the DESeq2 Wald statistic (*stat*). Global concordance was further validated by Pearson correlation of LFC values for significant DEGs, visualized as a four-quadrant scatter plot with linear regression. Pearson correlation coefficients (*R*) and linear regression lines were calculated using the ggpubr ^86^ package.

### Single-cell RNA-seq data processing

Raw sequencing reads from scRNA-seq libraries were aligned to the GRCm38 (mm10) reference genome using CellRanger ^87^ v6.1.2 (10x Genomics, Pleasanton, CA). Sparse data matrices generated by the CellRanger pipeline were processed further in R 4.3.2 using the Seurat v5 ^88^. Quality control included the removal of cells with mitochondrial content >20% and identification of doublets using DoubletFinder ^89^. To correct for batch effects between samples, Harmony ^90^ was used. Post batch correction, log normalization with a scale factor of 10,000 was performed on the count matrix. PCA was conducted, followed by dimension reduction using Uniform Manifold Approximation and Projection (UMAP) based on the first 10 principal components (PCs). Clusters of cells were identified using Seurat’s default shared nearest neighbor (SNN) modularity optimization method. To calculate the abundance of the TE-like population, the number of cells belonging to the cluster 4 was divided by the total number of cells for each genotype.

### Trajectory and regulon analyses

Trajectory analysis was performed using Monocle3 ^91^ to elucidate the developmental course across three cell types in the single-cell RNA-sequencing dataset. Cells were colored by pseudotime scores for visualization on the UMAP projections. DEGs along the pseudotime were identified by fitting a regression model to each gene. Regulon analysis was conducted using pySCENIC ^92^ in Python and SCENIC v1.3.1 ^93^ in R for integration with Seurat objects. Raw count matrices and cell metadata were converted to loom format compatible with pySCENIC. The GRNBoost2 algorithm was used to define regulons ^92^, which were further refined by filtering targets with significant enrichment for corresponding transcription factor (TF) motifs. The cellular activity of these regulons was calculated using the AUCell function and integrated with Seurat for further analysis ^93^. Differential regulons for each cell type were identified using Seurat’s FindAllMarkers function with a logFC threshold of 0.3 ^88^. Regulon activities were scaled for heatmap plotting. TFs representing each regulon cluster were subjected to gene set enrichment analysis, specifically with the ‘org.Mm.eg.db’ database and ‘Biological Process’ ontology ^94, 95^.

### Benchmarking of mouse embryo scRNA-seq data

For the benchmarking of our scRNA-seq data to the *in vivo* embryo counterparts, published expression matrices of mouse embryos (GSE45719 ^51^, GSE84892 ^52^, GSE100597 ^53^, GSE123046 ^54^) were collected. Embryonic stages were obtained from the original studies. Data for cells beyond E4.5 in Mohammed et al. ^53^ and Nowotschin et al. ^54^ were excluded. For the SMART-seq data from Deng et al. ^51^, the expression profiles derived from embryonic lysates, adult fibroblast, and liver were excluded. In the 10x-sequenced dataset from Nowotschin et al. ^54^, cells with < 2000 detected genes, and >10% mitochondrial content were excluded. Due to the large cell count compared to the other datasets, we aggregated the raw gene expression matrix of this dataset based on the neighborhood computation by miloR ^96^, instead of performing random downsampling. The neighborhood-based aggregation preserves the biological variation of the original data and mitigate sparsity and low sequencing depth, which are intrinsic to 10x single-cell datasets ^97^. Since the gene symbol formats used in the matrices were heterogeneous among datasets, gene symbols were first converted into Ensembl IDs based on the reference genome version used for alignment in each study. After merging, these Ensembl IDs were converted back to gene symbols using GRCm38 (Ensembl 2020 ver.) ^98^. Genes expressed in fewer than 3 cells were removed. The workflow from batchelor R package was implemented for integration ^99^. Per-batch scaling normalization was performed on the raw expression values of the merged dataset using multiBatchNorm function^99^. Dimensionality reduction was performed with 2,500 highly variable genes identified with Seurat’s SelectIntegrationFeatures function^100^, followed by MNN-based batch correction with fastMNN^99^ with the k-value of 15. 2D UMAP embedding was calculated with built-in umap function from uwot R package (https://CRAN.R-project.org/package=uwot) using 8 principal components. We identified clusters with Seurat’s FindNeighbors^100^ and FindClusters functions^100^. Then we annotated them using embryonic lineage marker expression profiles, after correcting log-expression values using batchelor’s mnnCorrect function ^99^. Trajectory analysis was performed with monocle3 ^101^. Query data projection was performed as previously described (Zhao *et al*. ^97^) with minor changes. For large (>500 cells) datasets, we used the same miloR aggregation approach ^96^ for down sampling. MNN pairs between query and reference were identified and filtered based on expression profile correlation. Corrected PC values of the query data were calculated based on the estimation of batch effects between MNN pairs. This approach ensures the positioning of the query data proximal to the cells in the reference cells with the most similar transcriptomic profiles. To compare WT, EPSC, and M2KO samples, the projection profiles of these samples were merged together with the reference embryos dataset. Published *in vitro* EPSC and TSC datasets were included as controls (Posfai *et al*.^102^, Angelova *et al*.^56^, Aldeguer *et al*. (https://doi.org/10.1101/510362)). Pseudotime assignment was performed over the *in vivo* Zygote-EPI and Zygote-TE branches using monocle3 ^101^. Expression correlation distances between query and reference were calculated based on pseudo-bulk log-normalized expression values. Pairwise Pearson correlations, computed with built-in cor() function (R 4.2.0), were converted to distance metrics (1-correlation), and scaled via min-max normalization. For DEG analysis between 3 clusters, public *in vitro* datasets were excluded to focus on the variation within our experimental data. To minimize batch effects in DEG analysis between 3 clusters, WT sample matched in batch with other groups were used. Seurat’s FindAllMarkers function was used with ‘wilcox’ option, with the ‘Late ICM’ cluster downsampled to 100 cells^100^. Analyses were run with 100 permutations, to derive the average adjusted p-value and log2 fold change value. Genes with average adjusted p-value >=0.05 were discarded. Top 200 genes by average log2 fold change values were used for heatmap visualization, with scaled pseudo-bulk values. GO enrichment was performed using enrichGO function of clusterProfiler ^103^.

### Immunofluorescence staining of Blastoids

The fixed structures were permeabilized in 0.2% Triton X in PBS for 10-15 min at RT, followed by PBST (1% Tween 20 in PBS) wash x2. Then the structures were blocked in PBS-5%BSA for 1h at RT. Primary antibodies were diluted in 0.5% BSA in PBST were incubated with the structures for O/N, at 4°C. The structures were then washed twice in PBST (1% Tween 20 in PBS). Secondary antibodies (1:500) diluted in 0.5% BSA in PBST were incubated with the structures for 3h, at RT. Nuclei were counterstained with DAPI at 100 ng/ml. The structures were washed with PBST x2, then gently loaded onto a confocal dish using pipette tips coated with 1% BSA in PBS. The solution was aspirated as much as possible, to allow the structures to adhere to the confocal dish. Then PBS was added to prevent drying. The primary antibodies and dilution ratios used are listed as following: anti-CDX2(1:250, Abcam, ab76541), anti-CDX2(1:150, Biogenx, Mu392A), anti-CK8(1:250, DSHB, TROMA1), anti-FOXA2(1:250, Abcam, ab108422), anti-Gata4(1:200, Santa Cruz Biotechnology, sc-25310), anti-E-cadherin(1:300, Cell Signaling Technology, 3195s). The secondary antibodies and dilution ratios used are listed as following: Alexa Fluor 405 Donkey anti-Rabbit IgG (H+L) (Invitrogen, A-48258), Alexa Fluor 647 Donkey anti-Rabbit IgG (H+L) (Invitrogen, A-31573), Alexa Fluor 647 Donkey anti-Mouse IgG (H+L) (Invitrogen, A-31571), Alexa Fluor 647 Goat anti-Rat IgG (H+L) (Invitrogen, A-21247).

### Immunostaining of Trophoblast Stem Cells

Round glass non-coated coverslips (Deckgläser) were coated with 0.5% porcine gelatin before seeding. Cells were seeded on coverslips and adapted to 2i/LIF media or EPSCM for 2 more days. After 4 days of TSC partial induction, coverslips were fixed with 4% paraformaldehyde for 10 min at RT, followed by permeabilization in 0.1% Triton X in PBS for 10 min at RT. Fixed cells were blocked in 3% BSA in PBS 1hr at RT or O/N at 4°C. The solution was aspirated, cells were incubated with specific primary and secondary antibodies for each 1hr at RT. The primary antibodies and dilution ratios used are listed as following: anti-CK8 (1:250, Sigma-Aldrich, ZRT1399). Secondary antibodies and dilution ratios used are listed as following: Alexa Fluor 647 anti-Rat IgG (H+L) (1:200, Cell Signaling, 4418S). Fluorescence images were examined with a TCS 8 confocal microscope (LEICA).

### Teratoma Formation assay

For teratoma formation assays, 7.5 × 10^5 of WT, M2KO#2, M2KO#8, EPSC cells were harvested and injected both subcutaneously (S.C. injection) and intratesticularly (I.T. injection) into BALB/c nude mice. Mice were euthanized 6 weeks post-injection, and teratomas were harvested and fixed in 10% neutral-buffered formalin (Biosesang) at 4°C. Fixed tissues were embedded in paraffin. After sectioning, hematoxylin and eosin (H&E) staining were performed for histological analysis.

### Nuclei isolation for CUT&Tag sample

After growing the cells to 70-80% confluency, media was removed from cell plate by suction. Freshly prepared conditioned media containing 0.1% formaldehyde (Sigma-Aldrich) was added and incubated at room temperature for 1 min. Subsequently, 500 μl of 2.5M glycine (GoldBio) was added dropwise, and the plate was placed on a shaker for 3 min. Cells were washed with 1x PBS, then 0.1% Trypsin-EDTA (Gibco) was added, and incubated at 37°C, CO2 incubator for 3 min. Any remaining cells were detached by tapping the plate, and conditioned media with 4 volumes of trypsin was added to inhibit enzyme activity. Cells were dispersed using a pipette-aid, centrifuged at 600g for 3 min at room temperature, and the supernatant was removed. The pellet was resuspended in 5 ml of cold Nuclear Extraction (NE) Buffer (20 mM HEPES (GoldBio) pH 7.9, 10 mM KCl (Duksan), 0.1% Triton X-100 (Duksan), 20% Glycerol (Daejung), 0.5mM Spermidine (Sigma-Aldrich), Protease inhibitors) and incubated on ice for 10 min. Next, the sample was filtered through a 30 μm strainer (Miltenyi Biotechnology) to remove aggregates and centrifuged at 600g for 3 min at 4°C to remove supernatant. The nuclei sample was aliquoted into cryovials and frozen slowly in a deep freezer using a cell freezing container.

### CUT&Tag and library preparation

CUT&Tag was performed based on Epicypher’s protocol. First, Concanavalin A (ConA) Paramagnetic Beads (Epicypher) were activated to bind nuclei sample using Bead Activation Buffer (20mM HEPES pH 7.9, 10mM KCl, 1mM CaCl_2_ (Duksan), 1mM MnCl_2_ (Duksan)). Activated ConA beads were incubated with 50,000 nuclei for 10 min at room temperature. Next, primary antibody was added to the nuclei-bead mixture in cold Antibody150 Buffer (20mM HEPES pH 7.5, 150mM NaCl (Duksan), 0.5mM Spermidine, Protease inhibitors, 0.01% Digitonin (Sigma-Aldrich), 2mM EDTA pH 8.0 (Thermo Scientific)) and incubated using a nutator overnight at 4°C. After the primary antibody was removed, secondary antibody mixture was added to nuclei-bead mixture in cold Digitonin 150 Buffer (20mM, HEPES pH 7.5, 150mM NaCl, 0.5mM Spermidine, Protease inhibitors, 0.01% Digitonin) and incubated using nutator during 30 min at room temperature. The nuclei-bead mixture was washed with Digitonin 150 Buffer three times and Digitonin 300 Buffer (20mM, HEPES pH 7.5, 300mM NaCl, 0.5mM Spermidine, Protease inhibitors, 0.01% Digitonin) was added before adding CUTANA pAG-Tn5 (Epicypher). CUTANA pAG-Tn5 pre-loaded with Illumina adapters was added to reactions and incubated in Digitonin 300 Buffer to bind antibody-labelled chromatin on a nutator for 1hr at room temperature. The nuclei-bead mixture with CUTANA pAG-Tn5 was activated with Tagmentation Buffer (20mM HEPES pH 7.5, 300mM NaCl, 0.5mM Spermidine, Protease inhibitors, 0.01% Digitonin and 10mM MgCl_2_ (Duksan)) for 1hr at 37℃ thermocycler. After washing three times with Digitonin 300 Buffer, reactions were stopped with TAPS Buffer (10mM TAPS pH 8.5 (Sigma-Aldrich) and 0.2mM EDTA) and quenched with SDS Release Buffer (10mM TAPS pH 8.5, 0.1% SDS (Enzynomics)) for 1hr at 58℃ thermocycler. SDS was neutralized with SDS Quench Buffer (0.67% Triton X-100). Finally, non-hot start PCR amplification was performed directly on the reaction mixture using universal i5 (Macrogen) and barcoded i7 primers (Macrogen), CUTANA High Fidelity 2x PCR Master Mix (Epicypher), and the DNA was purified using 1.3x AMPureXP beads (Beckman coulter).

### CUT&Tag sequencing and data analysis

CUT&Tag libraries were performed library quality check using an TapeStation HS D5000 Screen Tape (Agilent Technologies) to assess fragment size and DNA quality before proceeding to Illumina sequencing. Samples that passed the quality check were proceeded to paired-end Illumina sequencing using NovaSeq6000 (Illumina). Raw reads were trimmed for adaptor sequences using Trimmomatic v0.38 ^81^ and aligned to the mm10 reference genome using Bowtie2 v2.5.1 ^104^. Duplicate reads were removed with Picard v0.118 (https://broadinstitute.github.io/picard/). Mapped reads overlapping blacklist regions were removed prior to downstream analyses. We combined the ENCODE mouse blacklist (https://github.com/Boyle-Lab/Blacklist) and a CUT&RUN/CUT&Tag–specific blacklist^105^ and filtered reads with bedtools^106^ intersect function. Blacklist-filtered BAMs were used for all subsequent steps. Filtered reads were used to generate bigwig files using deepTools^107^ bamCoverage with binSize set to 50, effectiveGenome size of 2,652,783,500 for RPGC normalization, and ignoreDuplicates set to TRUE. These bigwig files were used as input for deepTools computeMatrix and plotHeatmap. For gene-centric heatmaps, reference-point mode with referencePoint TSS option was used with -b 3000 and -a 3000, alongside a bedfile corresponding to mm10 total genomic regions with skipZeros set to TRUE. Peak calling was performed with MACS2 ^78^. H3K27me3 CUT&Tag data peak calling was conducted with the broad option, while Mtf2 and H3K4me3 CUT&Tag data were called using the narrow option with a -q value of 1e-2. For peak annotation, MACS2 peak files were supplied to ChIPseeker^108^. TSS windows were set to ±1.5 kb and bar plots were generated with plotAnnoBar. PCA was performed using DiffBind v3.12.0 in R ^109^ on the read-count matrix derived from the merged peak set for each antibody, with library-size normalization to correct for technical batch effects while preserving biological variation. The same DiffBind framework with the DESeq2 method was used to identify differential H3K27me3 and H3K4me3 peaks for the contrasts M2KO vs WT and EPSC vs WT (FDR < 0.1, |log2 fold change| > 1). Differential peaks were annotated with ChIPseeker (TSS window = ±1.5 kb), and a gene-level chromatin status was assigned by selecting, for each gene, the most significant peak (highest |Fold|, lowest FDR). For H3K27me3, all annotation categories except “Distal Intergenic” were retained to accommodate its broad distribution; for H3K4me3, only peaks annotated to “Promoter” regions were retained. Genes were classified as positive for H3K27me3 loss (Fold < −1) or H3K4me3 gain (Fold > 1) based on their representative peak.

To identify Mtf2-dependent loci, we first merged all Mtf2 peak sets across WT and M2KO replicates using bedtools^106^ merge function. Mean Mtf2 signal was computed per region for WT (rep1 and 2) and M2KO (M2KO#2 rep1, 2 and M2KO#8 rep1, 2) using deepTools multiBigwigSummary on the merged peak bedfile. The M2KO mean signal was calculated as a hierarchical mean: replicates were averaged within each clone, and the two clone means were then averaged to assign equal weight to both clones. Regions with significant Mtf2 binding loss in M2KO relative to WT—defined as log2((WT_signal + 1) / (M2KO_signal + 1)) ≥ 1.5—were retained as “Mtf2-dependent regions”.These “Mtf2-dependent regions” were then used as references for heatmaps of Mtf2, H3K27me3, and H3K4me3 profiles. For these plots, computeMatrix reference-point --referencePoint center -b 3000 -a 3000 --binSize 50 --missingDataAsZero was applied to the bigwig files of all samples, followed by plotHeatmap with default clustering. Mtf2 binding loss in EPSC relative to WT was defined analogously, using the union of WT and EPSC Mtf2 broad peaks as the universe and applying the same log2 fold-change threshold of ≥1.5. Mtf2-loss regions were intersected with mm10 gene-body coordinates (GENCODE vM23) using bedtools intersect, and overlapping genes were called as ‘MTF2 loss positive’ in the corresponding cell type.

To test whether specific chromatin features are enriched in M2KO-upregulated TSC-related genes, we defined four mutually exclusive gene sets within the expressed-gene universe (n = 18,198 genes with non-NA adjusted P value in the M2KO-vs-WT DESeq2 result): Set A, M2KO-up DEGs that are also TSC-up DEGs (n = 612); Set B, M2KO-up DEGs not in the TSC signature (n = 1,568); Set C, TSC-up DEGs not upregulated in M2KO (n = 2,003); Set D, all expressed genes. TSC-up DEGs (n = 2,615) were obtained from GSE159181 (ESC vs TSC). For each set and each chromatin feature (H3K27me3 loss, MTF2 loss, H3K4me3 gain), the fraction of feature-positive genes was computed, and statistical enrichment was assessed by Fisher’s exact test (one-sided “greater”) for Set A vs Set B / C / D, with Benjamini–Hochberg correction within each comparison. Two-sided Fisher’s exact test was used to compute odds ratios with 95% confidence intervals. To compare chromatin redistribution between M2KO and EPSC, the gene sets defined for each chromatin feature in the two cell types were tested for overlap by Fisher’s exact test against the same expressed-gene universe (n = 18,198). Joint distributions of chromatin features were visualized with the UpSet function in ComplexHeatmap, and pairwise overlaps with the VennDiagram^110^ package.

For locus-level visualization of CUT&Tag and RNA-seq signal at selected genes, RPGC-normalized bigwig files were imported using rtracklayer^111^, and per-bin signal was computed within a strand-aware window of TSS −1 kb to +3 kb (200 equal-width bins, smoothed with a 5-bin centered moving average using zoo::rollmean). All replicates within each group (WT: 2 replicates; EPSC: 2 replicates; M2KO: 4 replicates from clones #2 and #8) were averaged to produce a single trace per group × antibody, and the y-axis was scaled per antibody to the 99.5th percentile of all values across the three groups. RNA-seq panels show DESeq2 size-factor-normalized counts as mean ± standard deviation; gene models depict the longest transcript per gene from TxDb.Mmusculus.UCSC.mm10.knownGene. Panels were assembled using ggplot2 and patchwork.

### Re-analysis of public transcriptomic datasets following PKC inhibition

To investigate the effect of Protein Kinase C (PKC) inhibition on the transcriptomic landscape and PRC2-mediated epigenetic regulation across species, we re-analyzed publicly available RNA-seq datasets of human and mouse ESCs. Human naïve ESC data were retrieved from the Gene Expression Omnibus (GEO) under accession number GSE153212. This dataset comprises two hESC lines (H9 and WIBR3) cultured under primed conditions and under naïve conditions with or without the PKC inhibitor Gö6983, providing a comparative framework for assessing the effect of PKC inhibition (sample composition shown in Fig. S7E). For mouse ESCs, transcriptomic data from cells cultured in 2i/LIF or PKCi-containing media were obtained from a previously published study ^64^.

Raw read counts were normalized using the variance-stabilizing transformation (VST) implemented in the DESeq2 package. To emphasize biological differences across pluripotent states and to mitigate technical variation arising from different cell lines and sequencing platforms, batch effect correction was performed using the removeBatchEffect function in the limma package; the biological group (culture condition) was retained as the variable of interest while cell line and platform were treated as batch covariates. Principal component analysis was performed on the batch-corrected VST values using the top 500 most variable genes (Fig. S7A for the mouse dataset; Fig. S7F for the human dataset).

Expression levels of core PRC2 subunits—*EZH1* and *EZH2* in the human dataset, and *Mtf2* and *Ezh2* in the mouse dataset—were extracted from the normalized and batch-corrected counts. For the human dataset, *EZH1* and *EZH2* expression was compared across primed, (−)PKCi naïve, and (+)PKCi naïve conditions (Fig. S7G). For the mouse dataset, *Mtf2* and *Ezh2* expression was compared between 2i/LIF and PKCi-containing conditions (Fig. S7B). Statistical significance was assessed using pairwise t-tests for the human dataset (multiple comparisons among three conditions) and unpaired t-tests for the mouse dataset (two-condition comparison), as indicated in the corresponding figure legends.

### Statistical analysis

Statistical analysis between two groups were performed using Student’s *t*-test, and in case of three or more groups with single variable, one-way analysis of variance (ANOVA) followed by Bonferroni comparison test was conducted. Two-way ANOVA was performed for groups with two or more variables. Dunnett’s multiple comparison test was performed in case of comparing experimental groups with single control group. Significance was set as P < 0.05 (*), P < 0.01 (**), P < 0.001 (***), P < 0.0001 (****). Error bars represent mean ± s.d.

## ACKNOWLEDGEMENTS

We are especially grateful to Sofiane Hamidi for his technical guidance and Yi Zhang for her assistance with pilot experiments. We also deeply thank the SignAC center at ASHBi for instrument access. We thank Dr. Jia-Ray Yu and Dr. Spyros Goulas for critically reviewing the manuscript and providing valuable comments and discussions. This work was supported by the National Research Foundation of Korea (NRF) grant, funded by the Korean government through the Ministry of Science and ICT (MSIT) [NRF2021R1C1C1013220 and NRF2022R1A5A102641311 for C.-H.L and RS-2023-00218543 and RS-2024-00432867 for H-J.C].

## AUTHOR CONTRIBUTIONS

H.-J.C. C.-H.L. M.C. and C.A conceived the overall study design, led the experiments, and wrote the manuscript. S.-M.K. and Y.-J.C. mainly conducted the cellular and biochemical experiments and data analysis. S.-K.L. and H.K. performed CUT&Tag-seq, bulk RNA-seq and cellular experiments. S.L, J.-W. B. performed bulk RNA-seq, scRNA-seq, and CUT&Tag-seq data processing, data analysis, visualization, and wrote the specific part of the manuscript. K.-T.K. produced the initial preliminary data. Y.-W.W. performed cellular experiments.

## COMPETING INTERESTS

The authors declare that they have no known competing financial interests or personal relationships that could have appeared to influence the work reported in this paper.

## DATA AVAILABILITY

The scRNA-seq, bulk RNA-seq and CUT&Tag datasets generated in this study have been deposited in the Gene Expression Omnibus (GEO) under the accession codes GSE306591 (scRNA-seq of *in vitro* cell models), GSE306592 (bulk RNA-seq of *in vitro* cell models) and GSE306593 (CUT&Tag of in vitro cell models profiling H3K27me3, MTF2 and H3K4me3). Publicly available datasets reanalyzed in this study were downloaded as provided by Liu *et al.* (GEO: GSE73952^24^ and GSE70605^112^), Deng *et al.*^51^ (GEO: GSE45719), Posfai *et al.* (GEO: GSE84892^52^ and GSE145609^102^), Mohammed *et al*.^53^ (GEO: GSE100597), Nowotschin *et al.*^54^ (GEO: GSE123046), Chen *et al.*^55^ (GEO: GSE74155), Angelova *et al.*^56^ (ENA: PRJEB68188) and Frias-Aldeguer *et al.*^113^ (GEO: GSE127754). The mouse reference genome (mm10, NCBI build 10) was obtained from the Genome Reference Consortium (https://www.ncbi.nlm.nih.gov/datasets/genome/GCF_000001635.20/), the gene annotation file from GENCODE (https://www.gencodegenes.org/mouse/release_M10.html), the ENCODE mm10 blacklist from Amemiya *et al.*^114^ (https://github.com/Boyle-Lab/Blacklist/blob/master/lists/mm10-blacklist.v2.bed.gz), and the CUT&RUN-specific mm10 blacklist from Nordin *et al.*^105^ All other data supporting the findings of this study are available from the corresponding authors on reasonable request.

## LEGENDS TO EXTENDED DATA FIGURES

**Extended data Figure 1.**
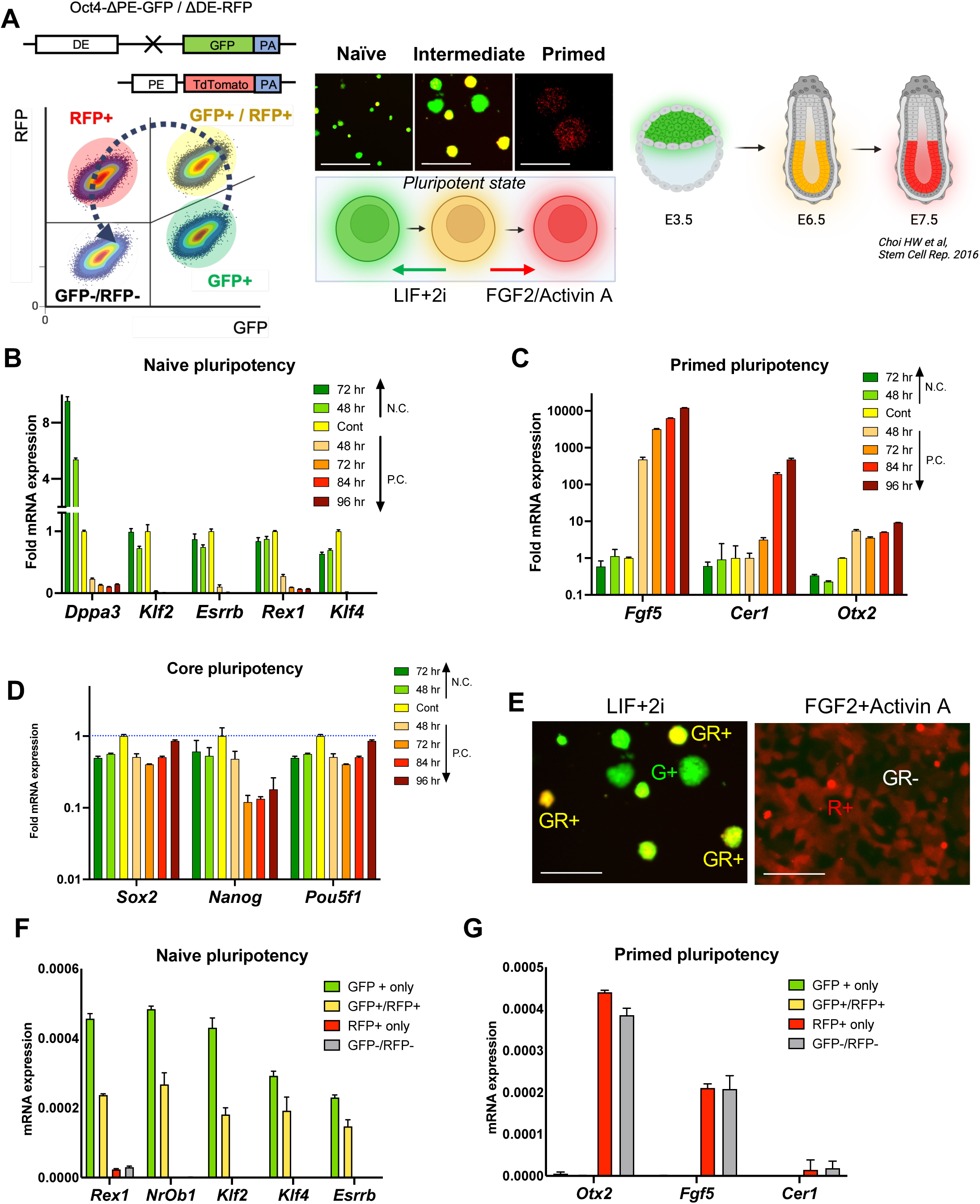
Dual reporter ESCs model for naïve and primed transition. (A) Graphical illustration of heterozygous Oct4-ΔPE-GFP allele, and Oct4-ΔDE-RFP allele (top left), flow cytometry data using OG2 GOF6 cell line (bottom left). Representative fluorescent images and graphical illustration of OG2 GOF6 cell line recapitulating naïve, intermediate, and primed pluripotent status (scale bars = 200 μm) (middle, right). [PE: Proximal Enhancer, DE: Distal Enhancer] (B) Relative mRNA expressions for naïve pluripotency marker genes (*Dppa3, Klf2, Esrrb, Rex1 and Klf4*) during either naïve or primed conversion, (n = 3). (C) Relative mRNA expressions for primed pluripotency marker genes (*Fgf5, Cer1, Otx2*) during either naïve or primed conversion, (n = 3). (D) Relative mRNA expressions for core pluripotency marker genes (*Sox2, Nanog, Pou5f1*) during either naïve or primed induction, (n = 3). (E) Representative fluorescence images for OG2 GOF6 cells cultured under serum/LIF+2i condition (Left) and Fgf2+ActivinA condition (right) (scale bars = 250 μm). (F) Relative mRNA expressions for naïve pluripotency marker genes (*Rex1, NrOb1, Klf2, Klf4, Esrrb*) of four indicative populations, (n = 3). (G) Relative mRNA expressions for primed pluripotency marker genes (*Otx2, Fgf5, Cer1*) of four indicative populations, (n = 3).

**Extended data Figure 2.**
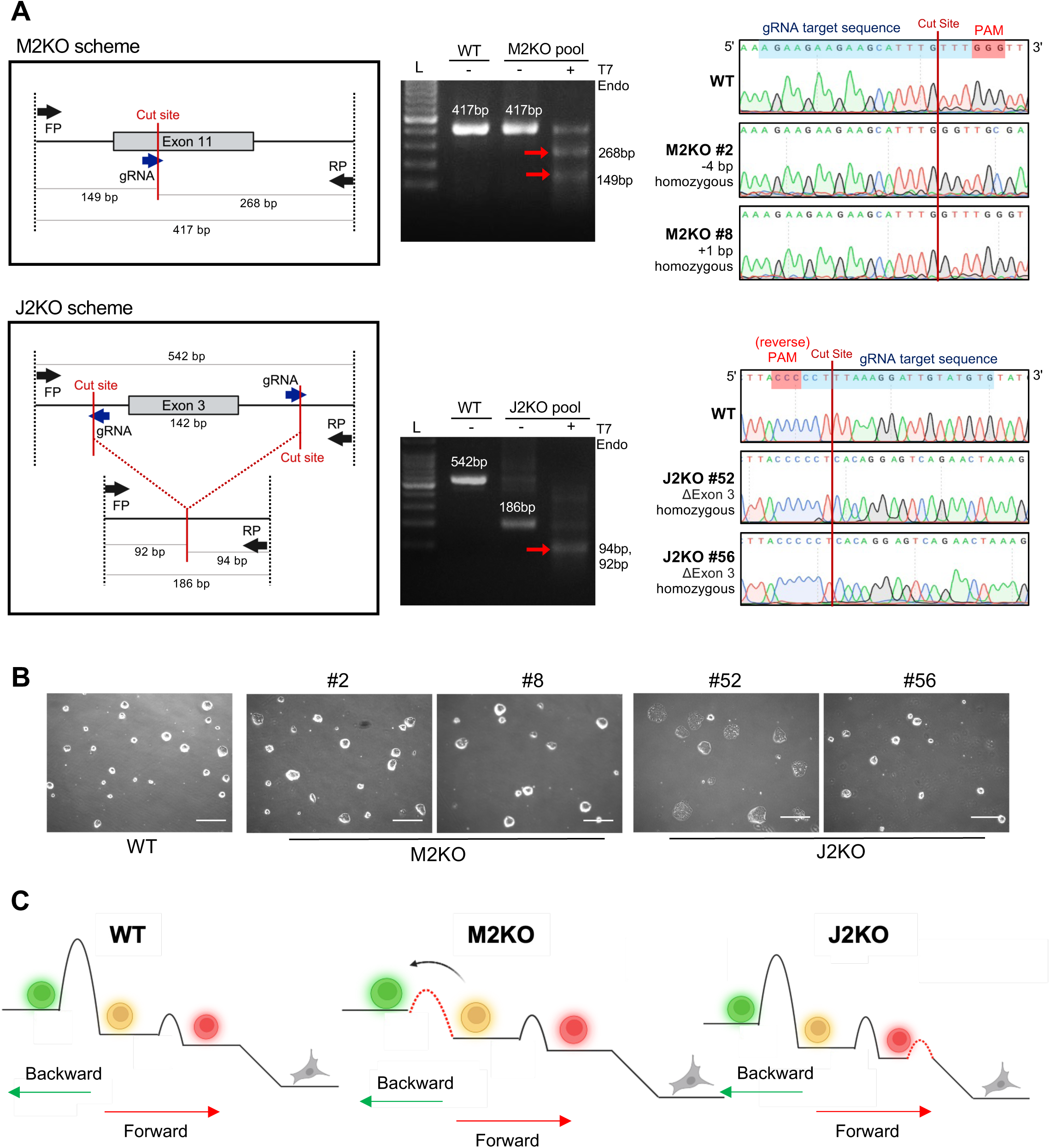
Establishment of M2KO and J2KO ESCs. (A) Graphical outline for generation and validation of Mtf2 KO (hereafter, M2KO) (top left). Graphical outline for generation and validation of Jarid2 KO (hereafter, J2KO), exon 3 (142 bp) was deleted using two gRNAs targeting the flanking intronic regions (bottom left). T7 Endonuclease I assay for WT and M2KO pool (top middle). Sanger sequencing chromatograms of WT and individual M2KO clones with PAM sequence highlighted (top right). T7 Endonuclease I assay for WT and J2 KO pool (bottom middle). Sanger sequencing chromatograms of WT and individual J2KO clones with complementary PAM sequence highlighted (bottom right). FP, forward primer; RP, reverse primer. (B) Representative brightfield images of OG2 GOF6 WT, M2KO#2, M2KO#8, J2KO#52, and J2KO#56 (scale bars = 500 μm). (C) Graphical illustration indicating epigenetic barriers from naïve pluripotency to differentiation in WT, M2KO and J2KO.

**Extended data Figure 3.**
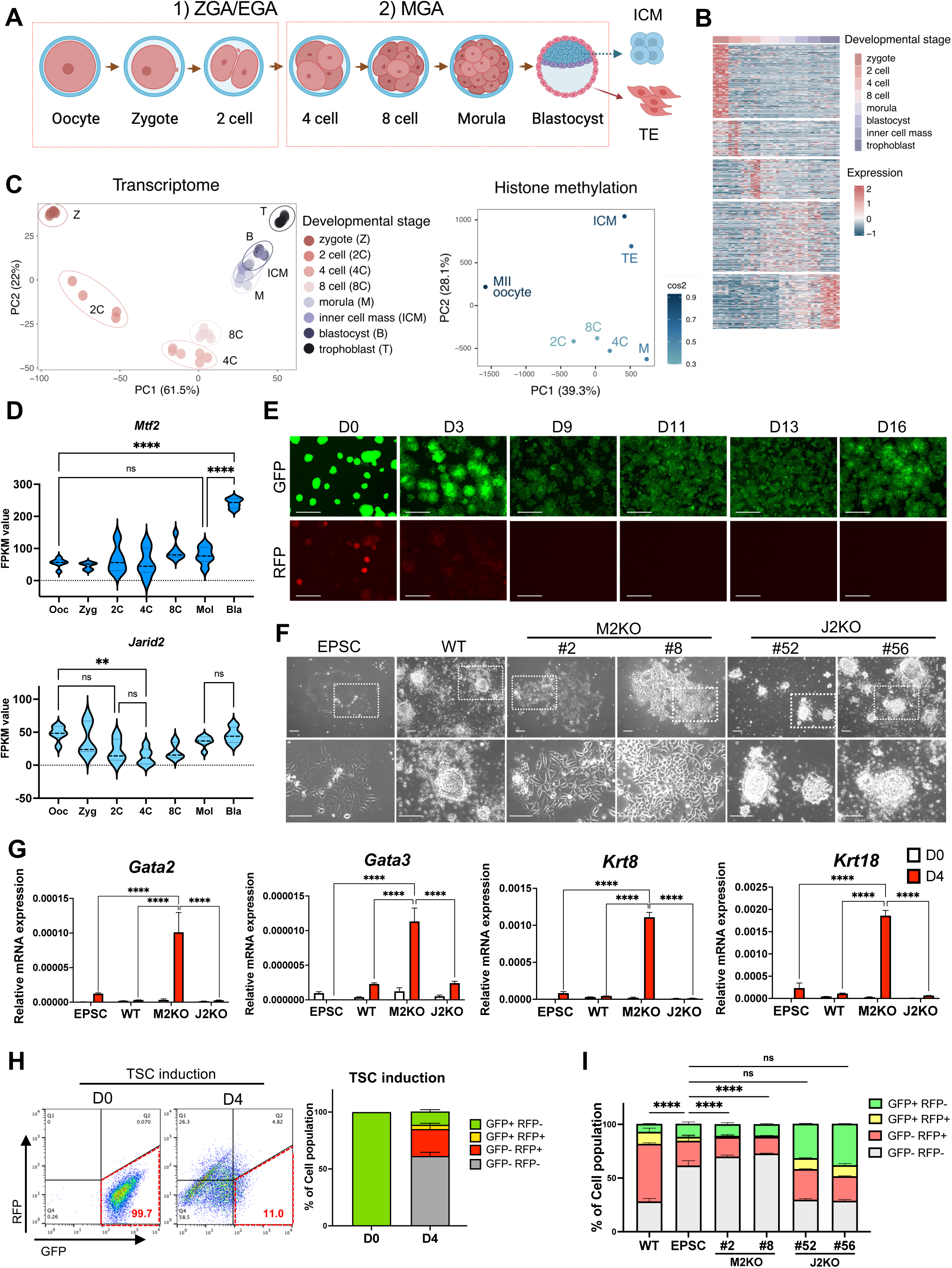
Differentiation to TSCs. (A) Graphical summary of embryonic developmental stages (B) Principal Component Analysis (PCA) of RNA-seq data across eight distinct embryonic developmental stages (C) PCA of ChIP-seq data across seven different stages, emphasizing the bivalency pattern of H3K27me3 and H3K4me3 profiles. The color bar in the right panel indicates the quality of representation for each variable on the factor map, determined by squared cosine. (D) Violin plot of FPKM values of indicative genes (*Mtf2* and *Jarid2*) at indicative stages of embryo (n = 4, 5, 6) (**, p < 0.01, ****, p < 0.0001, n.s. for non-significant). (E) Relative fluorescence images at indicative days during EPSC induction (scale bars = 500 μm). (F) Representative bright field images of EPSC, WT, M2KO#2, M2KO#8, J2KO#52 and J2KO#56 (scale bars = 200 μm, top panels, scale bars = 50 μm, bottom panels) after 4 days of partial TSC induction. (G) Relative mRNA expressions for TSC marker genes (*Gata2*, *Gata3*, *Krt8*, *Krt18*) in EPSC, WT, M2KO, and J2KO at Day 0 and Day 4 of partial TSC induction. Two-way ANOVA, multiple comparisons (****, p < 0.0001), (N=1, n = 3). (H) Flow cytometry data of EPSCs and TSCs (left), quantification of each population (right), (N = 1, n = 3). (I) Quantification of each indicative cell population of flowcytometry data at day4 of TSC partial induction (****, p < 0.0001, n.s. for non-significant), (N = 1, n = 3).

**Extended data Figure 4.**
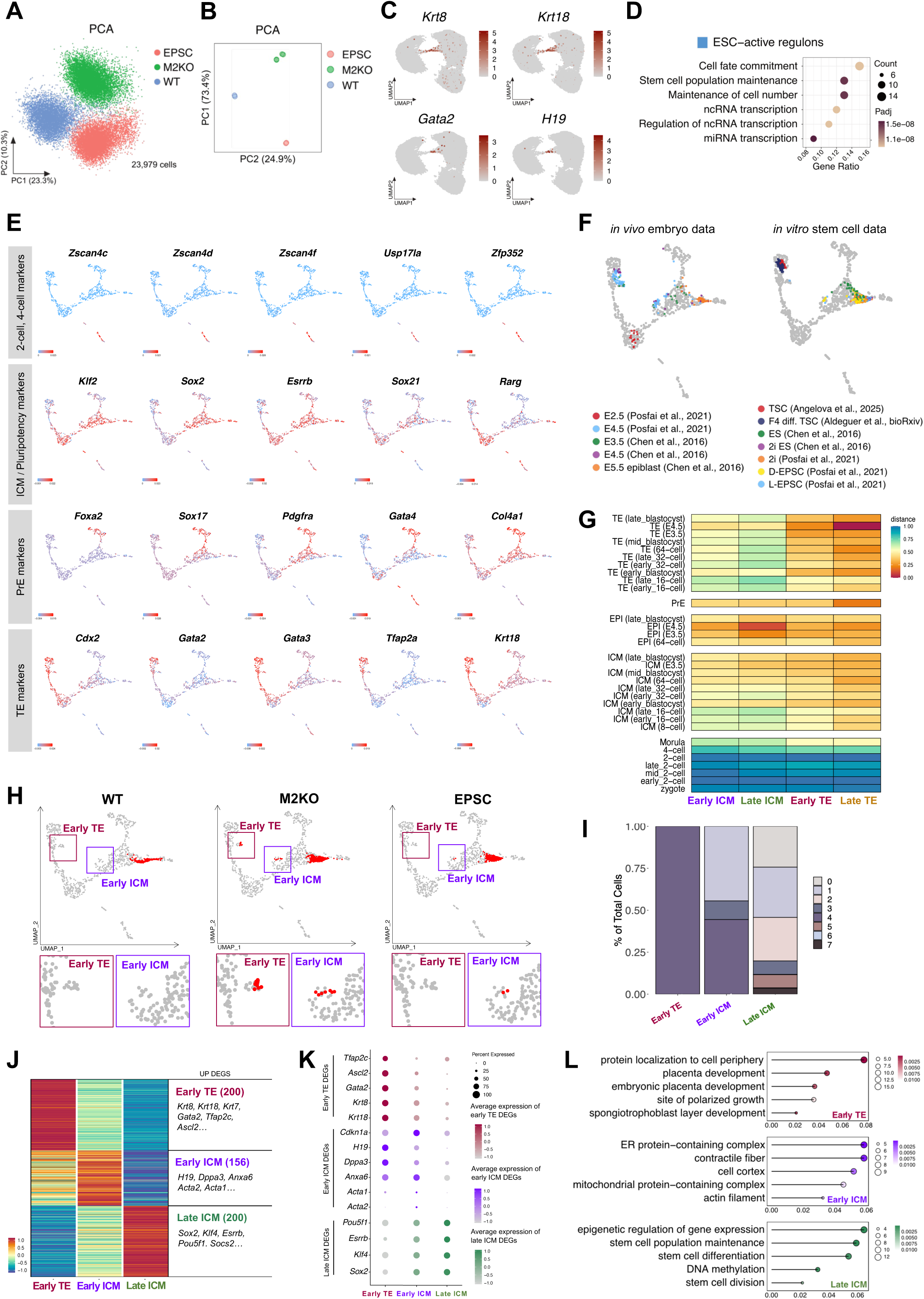
Transcriptome analysis. (A) PCA of scRNA-seq data. (B) PCA of bulk RNA-seq data. (C) UMAP feature plots showing the normalized expression of representative trophectoderm (TE) markers (*Krt8, Krt18, Gata2*) and the totipotency-associated marker (*H19*) in the experimental single-cell dataset, highlighting their localized and specific enrichment within cluster 4. (D) GO terms significantly enriched (*P*_adj_ < 0.05) for TFs comprising “ESC-active” regulon clusters, identified by gene set enrichment test. (E) UMAP plot of integrated scRNA-seq data from *in vivo* mouse embryos, showing the expression of key marker genes for early embryonic lineages. (F) UMAP projection of public single-cell datasets for *in vivo* embryo data (left) and *in vitro* stem cell data (right), onto the integrated embryonic reference. (G) Heatmap of pairwise correlation distance metrics between transcriptomic profiles of *in vitro* clusters and *in vivo* embryonic data stratified by embryonic stage and lineage. (H) UMAP projection of WT, M2KO, and EPSC cells mapped onto the integrated in vivo embryo data, with projected cells highlighted in red. Rectangles on the bottom provide zoomed-in views of cells localized within the early TE and early ICM clusters. (I) Bar plot showing the proportion of clusters from Extended data Fig.4C mapped to each identified cluster. (J) Differential gene expression analysis between clusters within experimental dataset. (K) Dot plot with representative differentially expressed genes (DEGs) for each cluster. (L) Five enriched Gene ontology (GO) terms associated with the DEGs from (K) are displayed.

**Extended data Figure 5.**
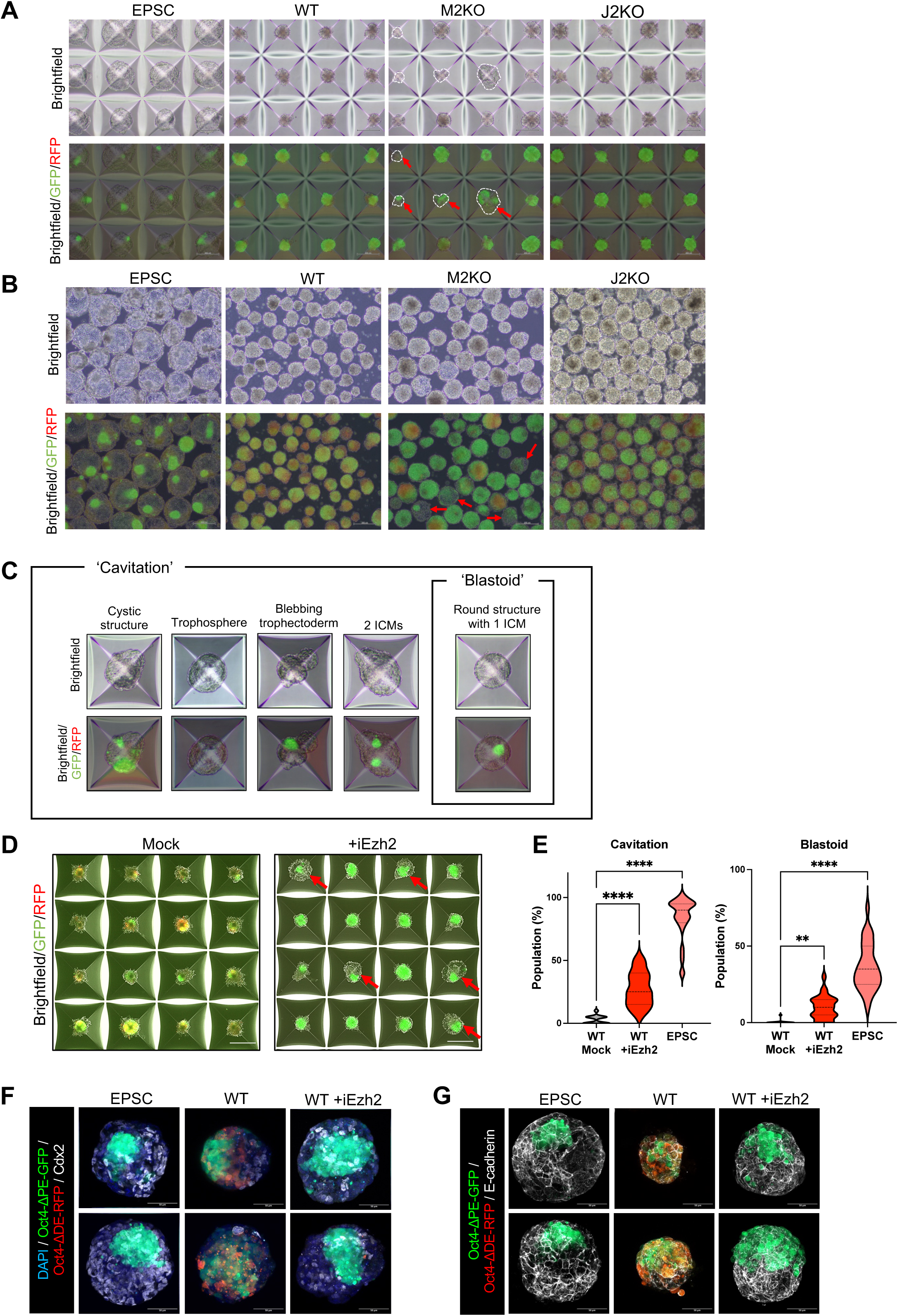
Establishment of blastoid. (A) Representative brightfield and brightfield/GFP/RFP images of structures derived from EPSC, WT, M2KO, and J2KO on blastoid formation day 7, on AggreWell. White arrows indicate cavitations of M2KO (scale bars = 200 μm). (B) Representative Brightfield and brightfield/GFP/RFP images of structures derived from EPSC, WT, M2KO, and J2KO on blastoid formation day 7, harvested from AggreWell. Red arrows indicate cavitations of M2KO (scale bars = 200 μm). (C) Representative brightfield and brightfield/GFP/RFP images of criteria for quantification of structures. (D) Representative brightfield/GFP/ RFP images of EPSC and WT with or without iEzh2 treatment, on day 7 of blastoid formation (scale bars = 200 μm). (E) Quantification of ‘Cavitation’ and ‘Blastoid’ structures derived from EPSC, WT without iEzh2, and WT with iEzh2 on blastoid formation day 7. Total 535 structures of WT mock, 535 structures of WT +iEzh2, and 617 structures of EPSC were analyzed. Each point represents the percentage of ‘Cavitation’ and ‘Blastoid’ structures in a single imaged field (27 fields for each of WT mock and WT+iEzh2, and 31 fields for EPSC). Mixed-effects analysis, Multiple comparisons, (****, p < 0.0001, **, P < 0.01, N=3, n=27, 31). (F-G) Immunofluorescence staining of structures derived from EPSC, WT without iEzh2, WT with iEzh2 on day 7 of blastoid formation for Cdx2 and DAPI (F) and for E-cadherin (G) (scale bars = 50 μm).

**Extended data Figure 6.**
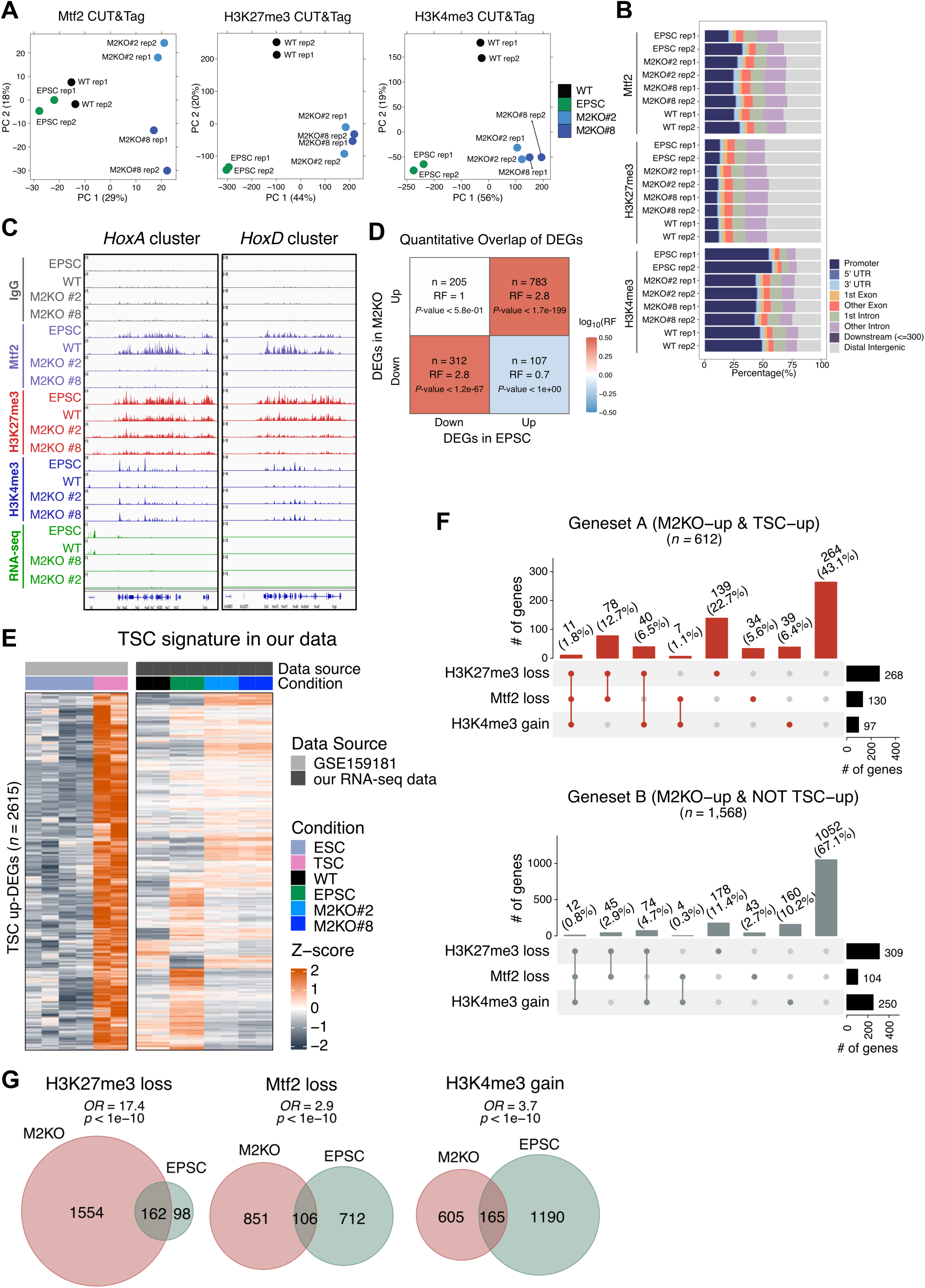
Integrated transcriptomic and epigenomic analyses. (A) PCA of CUT&Tag peak profiles for Mtf2, H3K27me3, and H3K4me3 in four different conditions. For Mtf2 (left), PCA was performed specifically using Mtf2-dependent signals; for H3K27me3 (middle) and H3K4me3 (right), PCA was performed using global peak profiles. (B) Genomic distribution of Mtf2, H3K27me3, and H3K4me3 peaks across promoters, UTRs, exons, introns, downstream (≤300 bp), and distal intergenic regions. (C) Mtf2, H3K27me3 and H3K4me3 CUT&Tag and RNA-seq genome browser tracks of *HoxA* (left) and *HoxD* (right) clusters in 4 different conditions. (D) Quantitative overlap analysis of differentially expressed genes (DEGs) between EPSC vs. WT and M2KO vs. WT. The number of overlapping genes (n), Representation Factor (RF), and hypergeometric *P*-values are indicated. Color scale represents log_10_(RF). (E) Heatmap of TSC-signature genes (n=2,615 TSC-up DEGs from GSE159181, ESC vs TSC) in the public dataset (left) and our RNA-seq data (right). Expression is shown as Z-scored counts. (F) Upset plots showing joint distribution of three chromatin features (H3K27me3 loss, Mtf2 loss, and H3K4me3 gain) within Set A (top, M2KO-up & TSC-up; n=612) and Set B (bottom, M2KO-up & not TSC-up; n=1,568). Vertical bars indicate the number and percentage of genes in each combination; right horizontal bars indicate marginal totals per mark. (G) Mark-wise overlap of differentially affected genes between M2KO and EPSC for each chromatin feature: H3K27me3 loss (left), Mtf2 loss (middle), and H3K4me3 gain (right). Each feature was defined relative to WT in the corresponding cell type. Odds ratios and one-sided Fisher’s exact-test *P* values were calculated against the universe of expressed genes (n=18,198).

**Extended data Figure 7.**
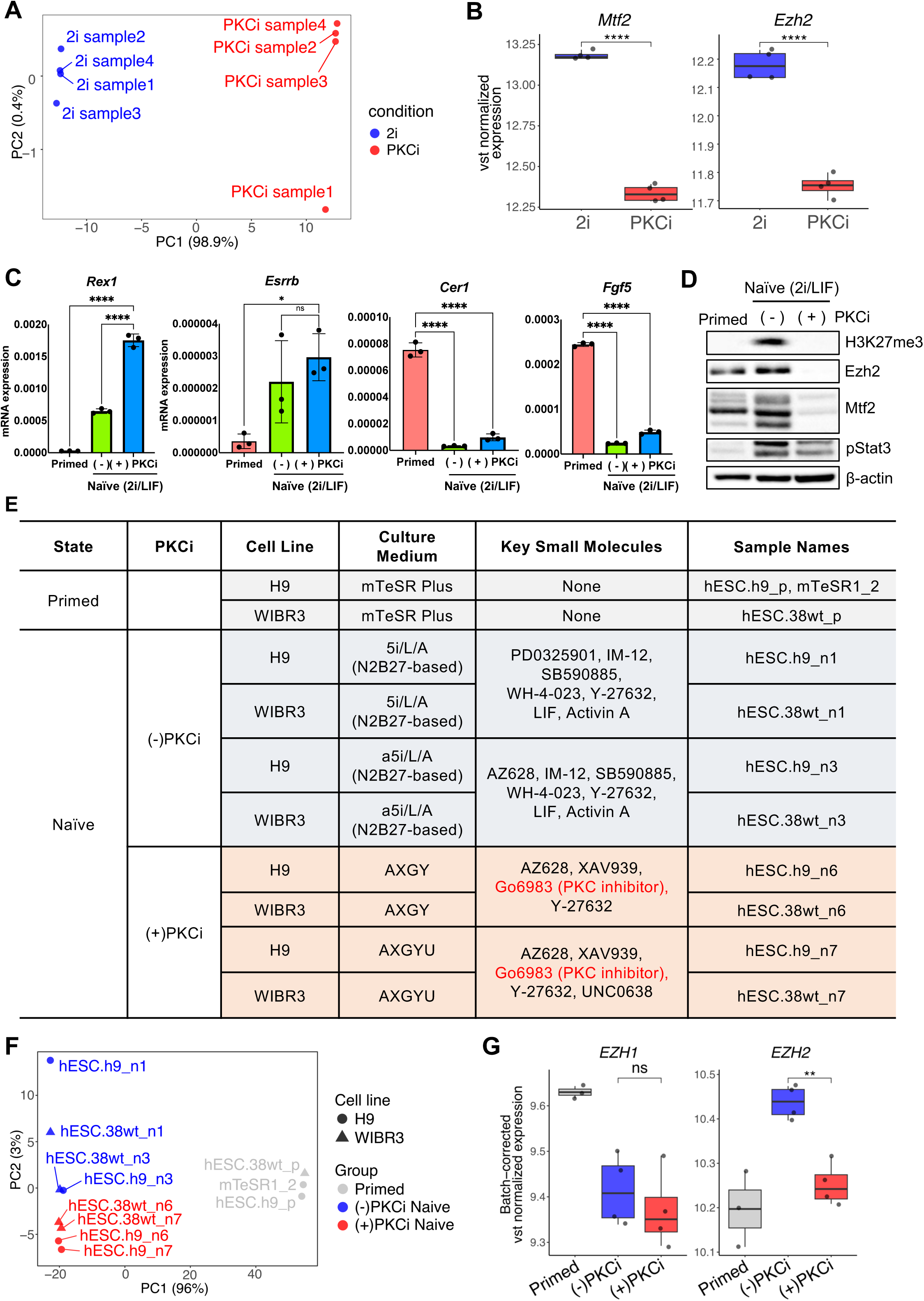
PKC inhibition attenuates Mtf2-PRC2 activity in naïve pluripotency. (A) Principal component analysis of bulk RNA-seq from a published mouse ESC dataset. Cells cultured under conventional 2i/LIF (blue) and Gö6983-containing PKCi conditions (red) form transcriptionally distinct clusters along PC1 (98.9% of variance). (B) *Mtf2* and *Ezh2* expression in the same dataset, shown as variance-stabilizing transformed (VST) normalized counts. Both genes are significantly downregulated upon PKCi treatment (unpaired *t*-test; ****, *P* < 0.0001). (C) mRNA expressions for naïve pluripotency marker genes (*Rex1, Esrrb*) and primed pluripotency marker genes (*Cer1, Fgf5*) of primed, naïve and naïve-converted primed with Gö6983 (2.5µM) under serum LIF+2i condition, One-way ANOVA, multiple comparisons, (*, p < 0.05, ****, p < 0.0001, n.s. for non-significant). (N = 1, n = 3). (D) Immunoblotting analysis for indicative proteins (H3K27me3, Ezh2, Mtf2, pStat3) in primed, naïve and naïve-converted primed mESC with PKC inhibitor Gö 6983(2.5µM). β –actin was used as a loading control. (E) Sample information for the human naïve ESC bulk RNA-seq dataset (GSE153212). Samples cover two cell lines (H9, WIBR3) and three pluripotent states: primed, (−)PKCi naïve, and (+)PKCi naïve. Key small-molecule supplements are listed for each condition. (F) Principal component analysis of the human naïve ESC dataset in (E), performed on the top 500 most variable genes after VST and limma-based batch correction across cell lines and platforms. Samples are colored by culture condition (primed: gray; (−)PKCi naïve: blue; (+)PKCi naïve: red) and shaped by cell line (circle: H9; triangle: WIBR3). (G) *EZH1* and *EZH2* expression in the human naïve ESC dataset, shown as VST-normalized and batch-corrected counts. *EZH2* is significantly downregulated in (+)PKCi naïve hESCs relative to (−)PKCi naïve, whereas *EZH1* is comparable. Pairwise *t*-tests (**, P < 0.05, ns, non significant).

## Notes

### Competing Interest Statement

The authors have declared no competing interest.

