## Supplementary figures for "Mtf2 Safeguards Naïve Pluripotency by Restricting Trophectoderm Competence"

### Extended data Figure 1

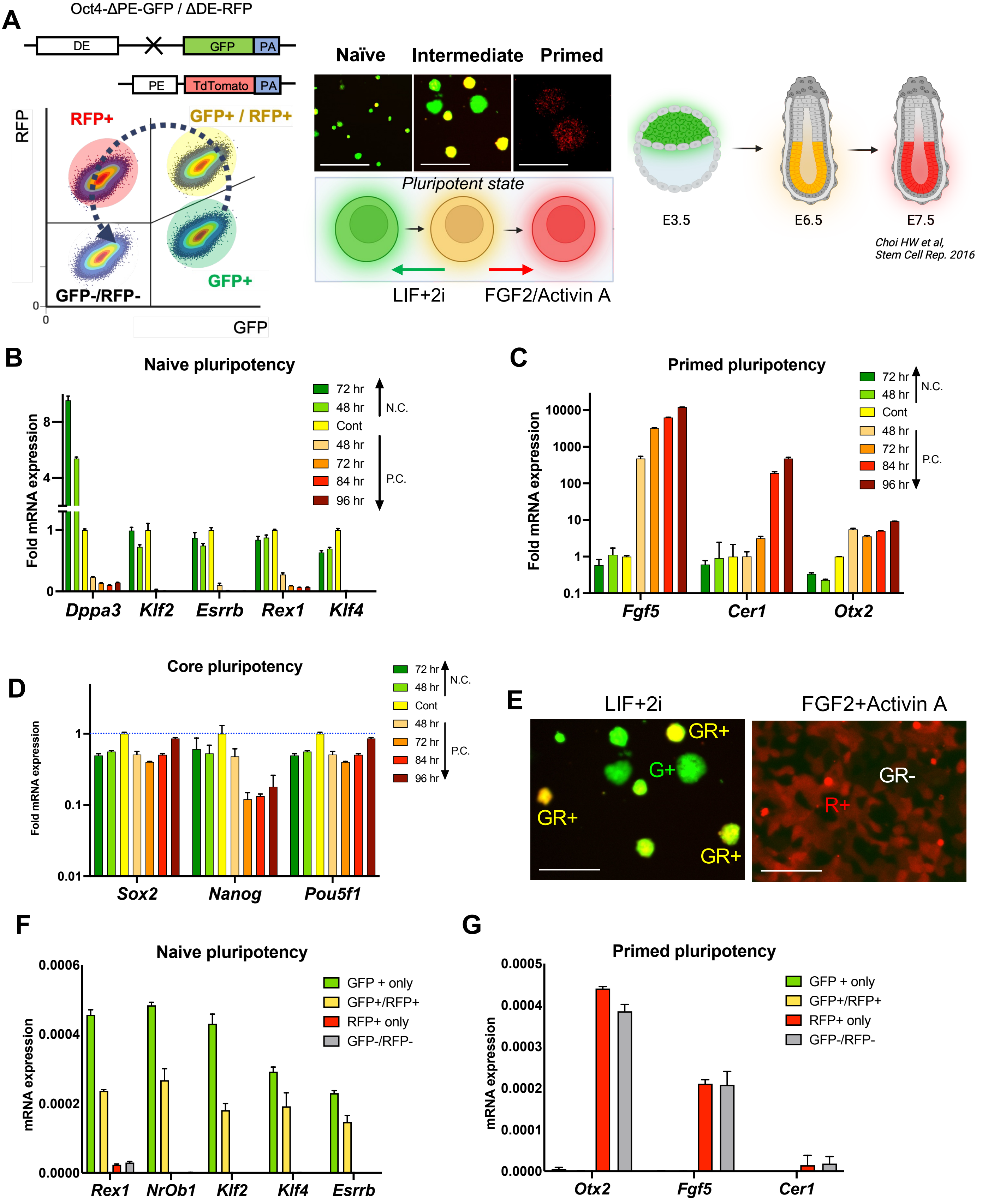

### Extended data Figure 2

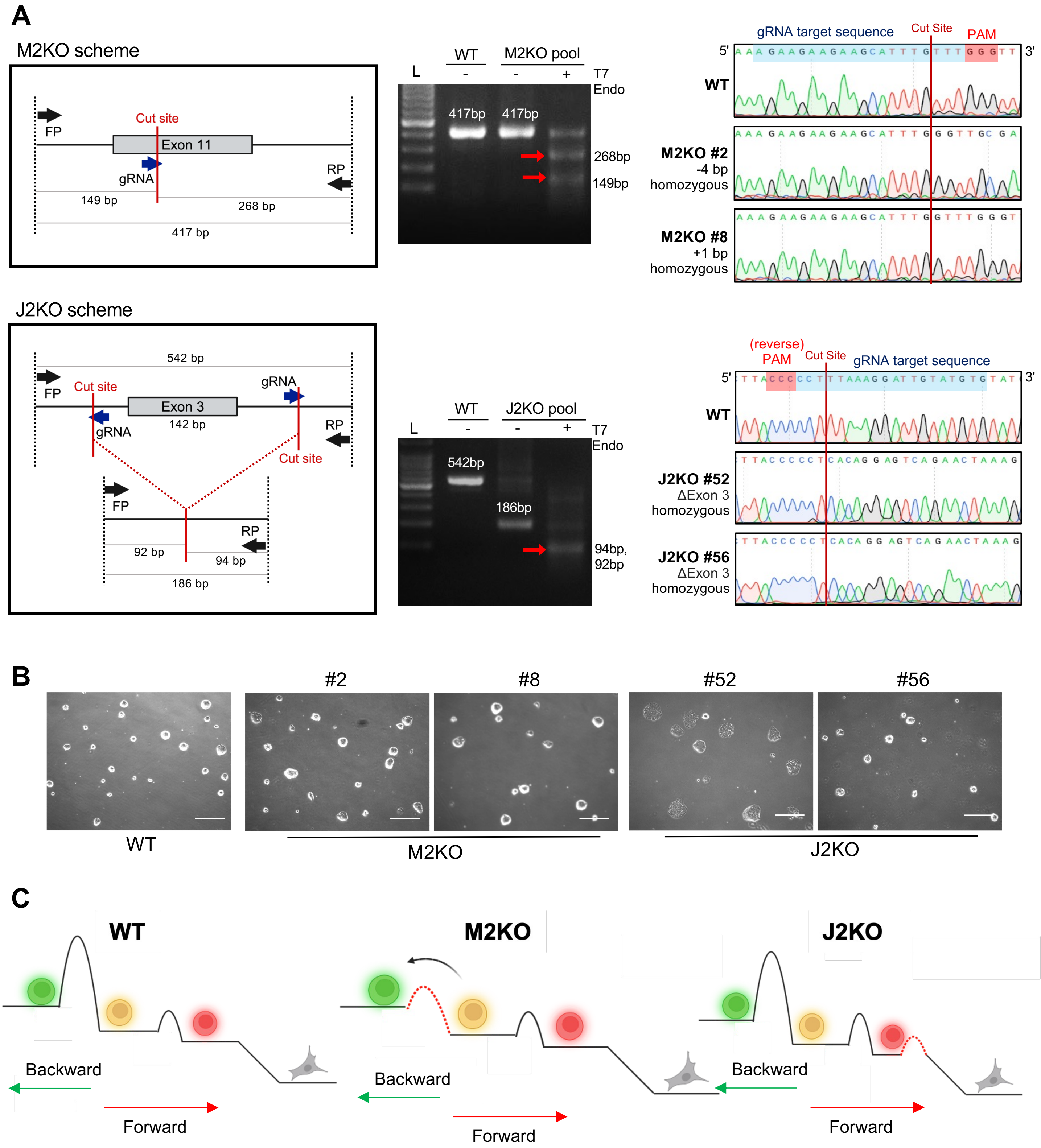

### Extended data Figure 3

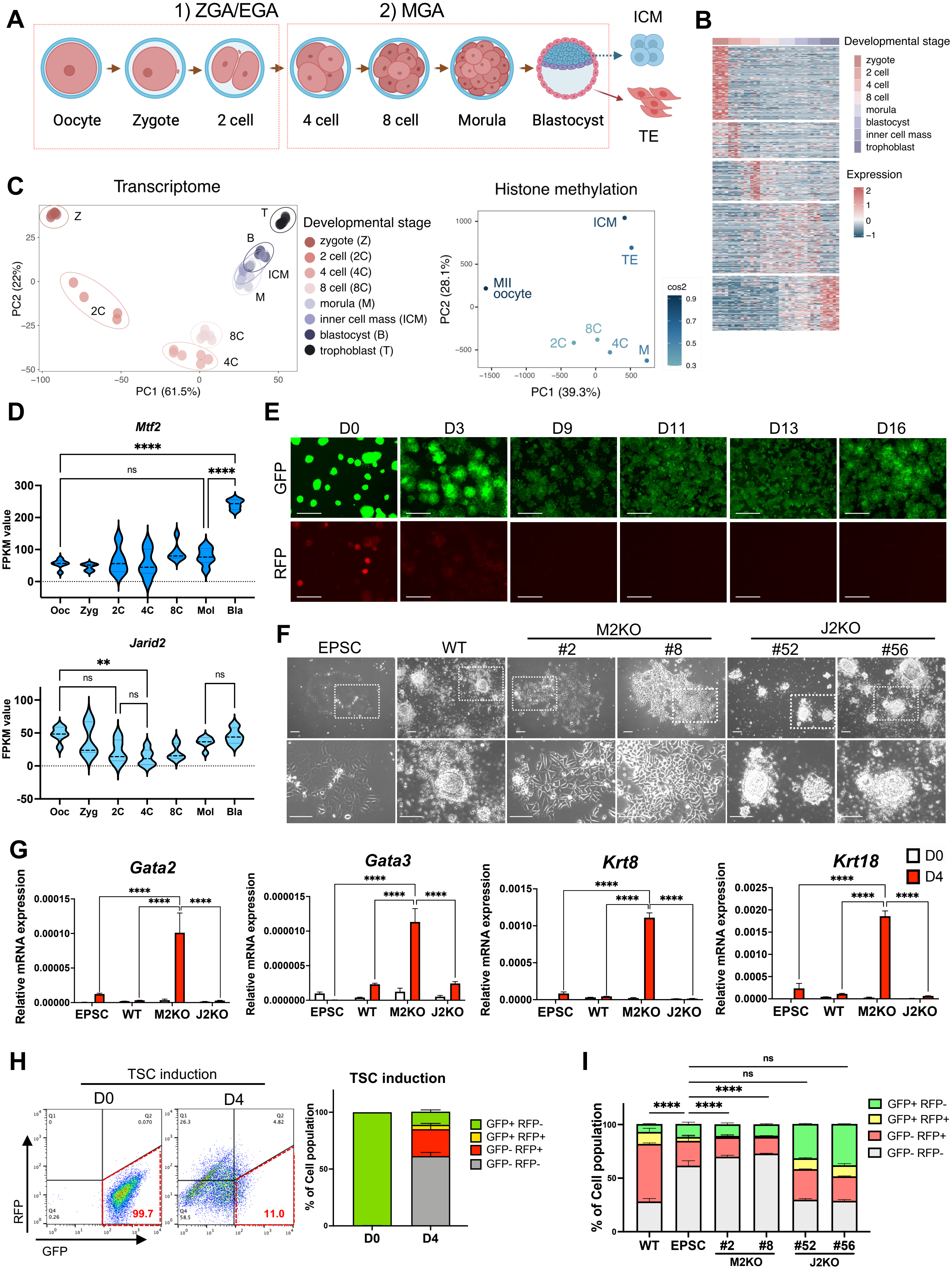

### Extended data Figure 4

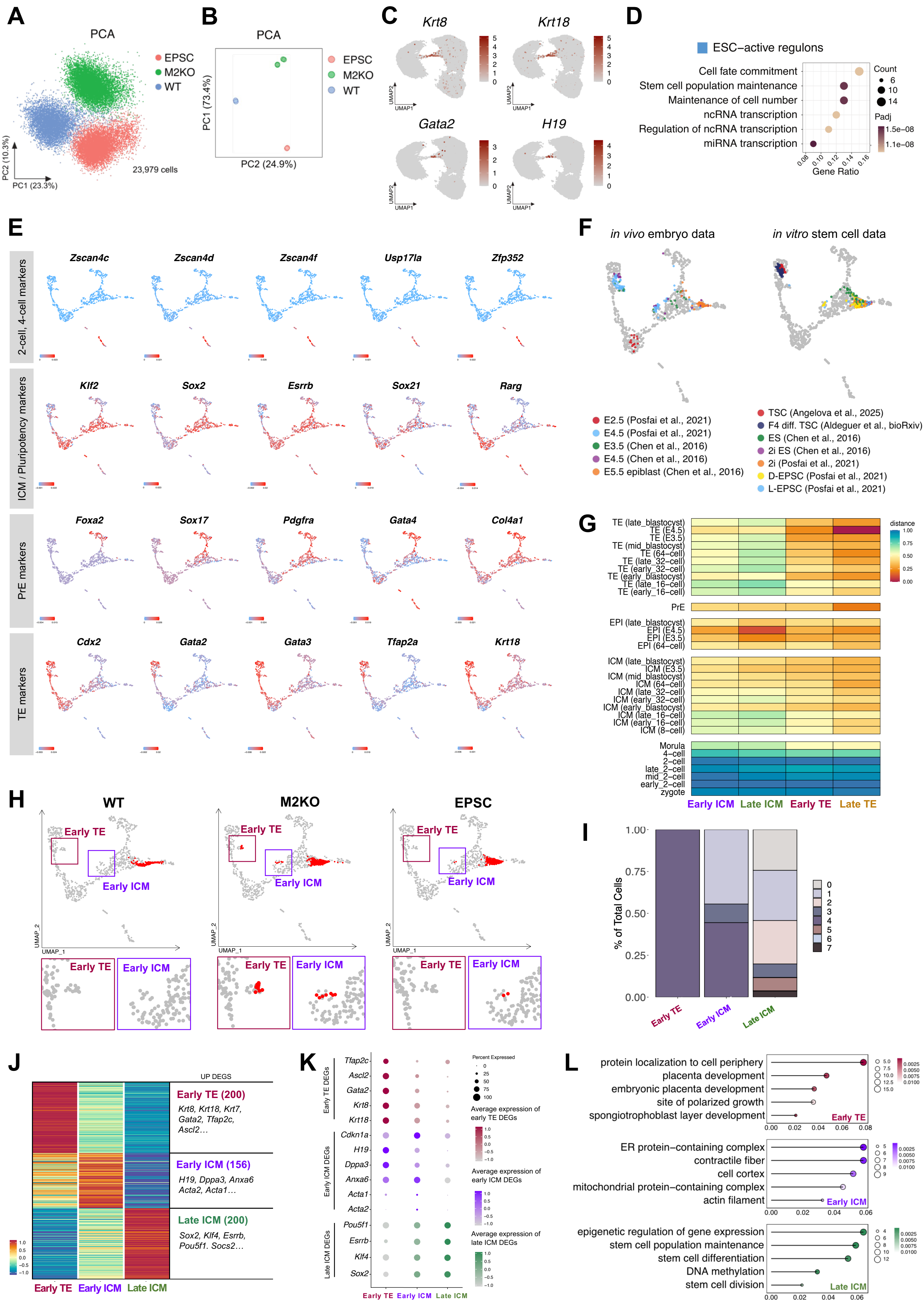

### Extended data Figure 5

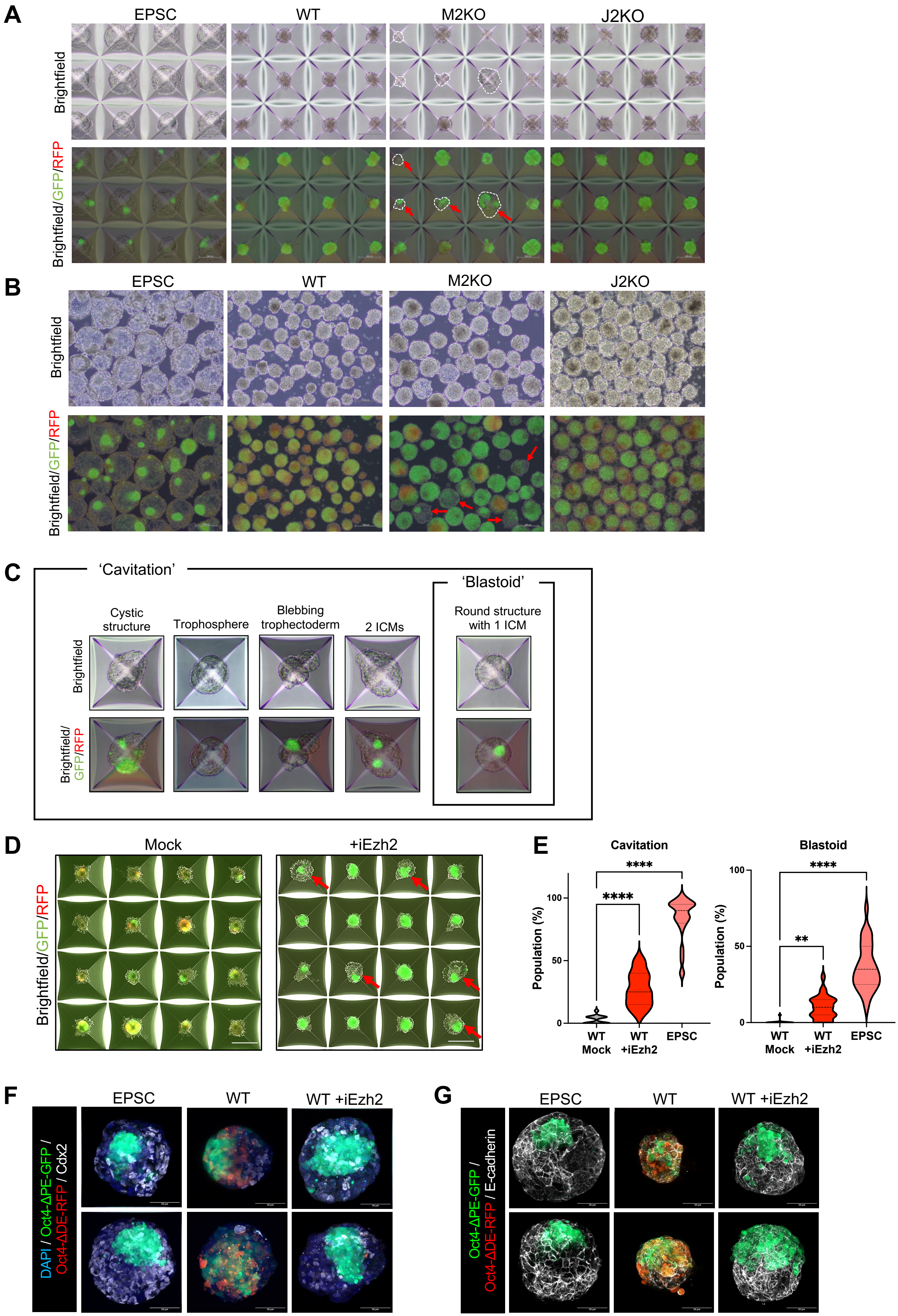

### Extended data Figure 6

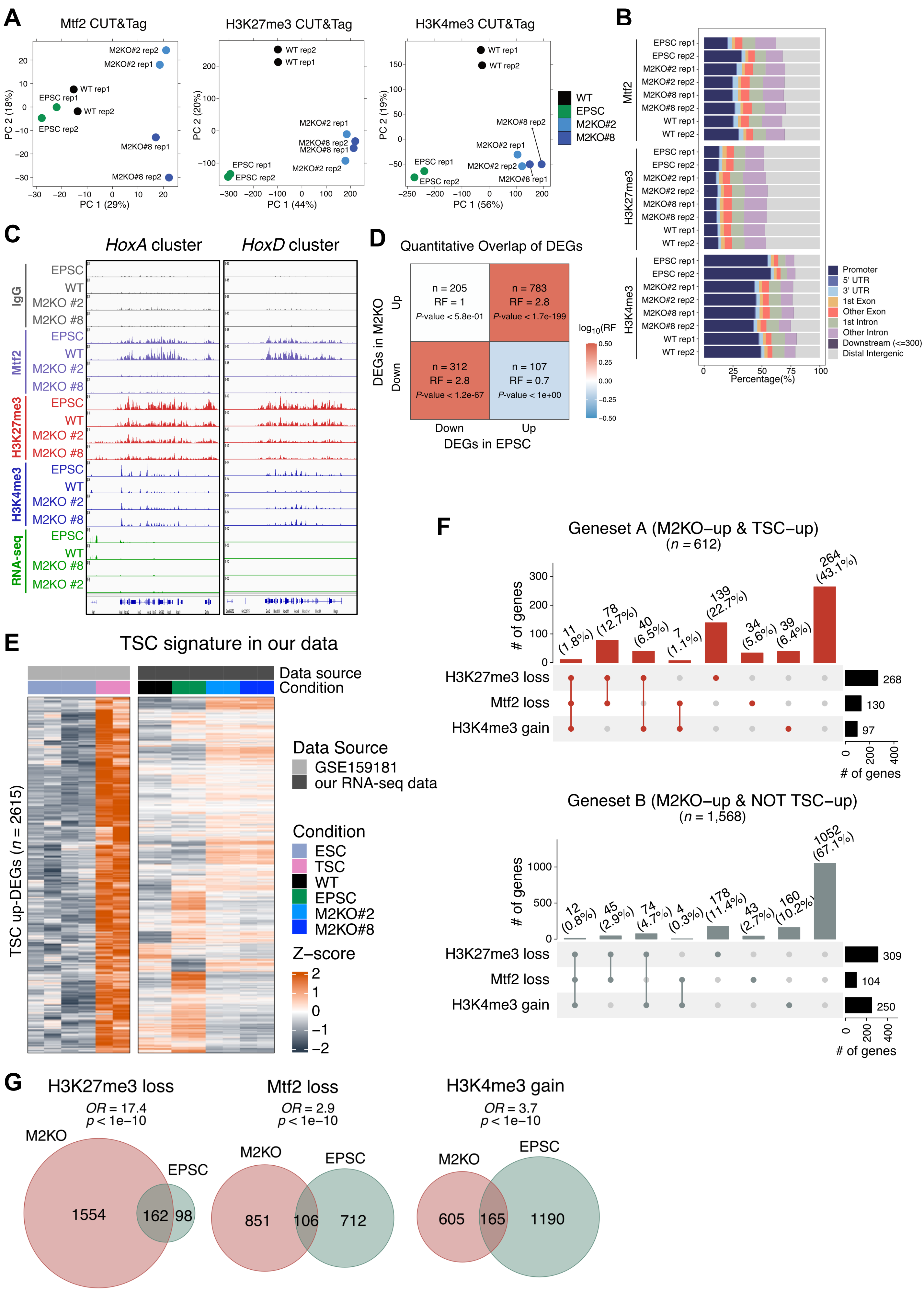

### Extended data Figure 7

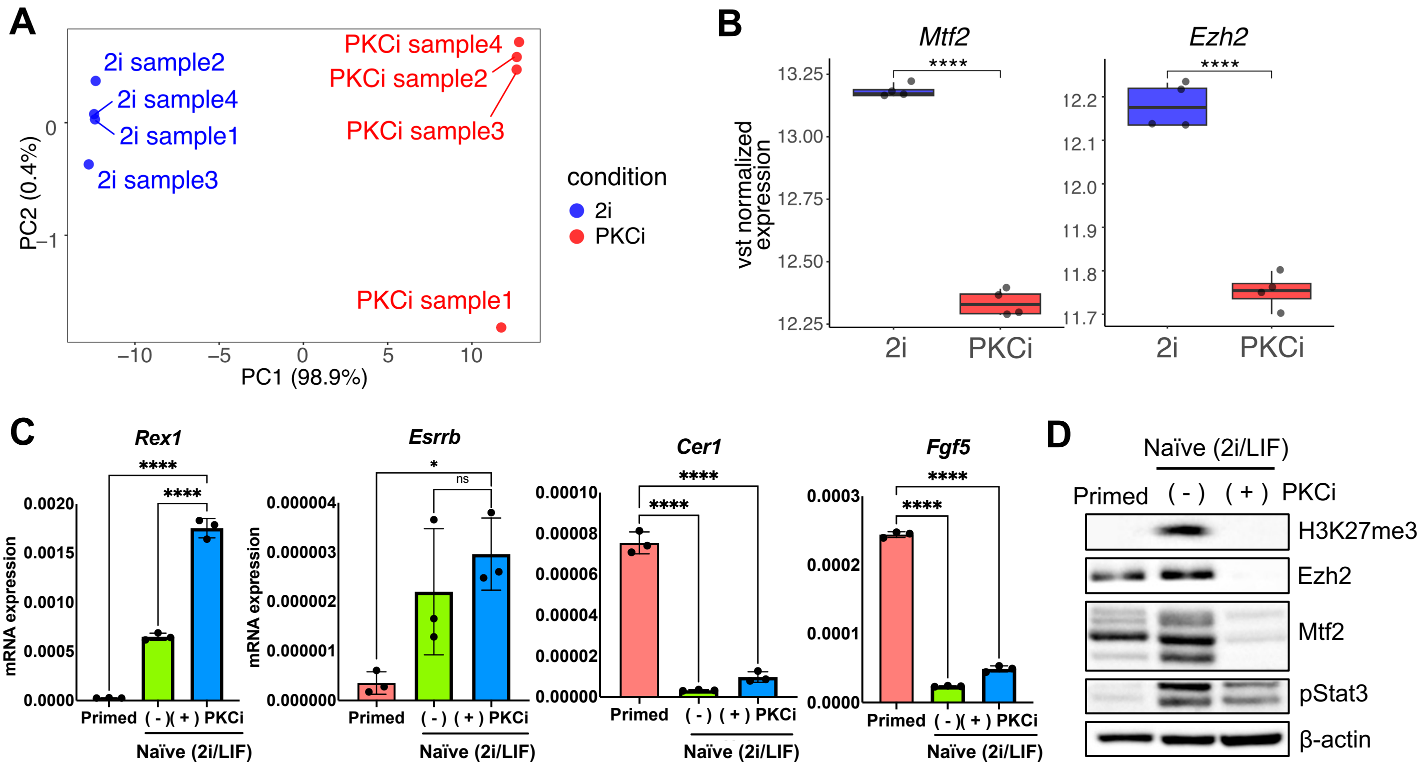


E

| State | PKCi | Cell Line | Culture Medium | Key Small Molecules | Sample Names |
|---|---|---|---|---|---|
| Primed | | H9 | mTeSR Plus | None | hESC.h9_p, mTeSR1_2 |
| | | WIBR3 | mTeSR Plus | None | hESC.38wt_p |
| Naïve | (-)PKCi | H9 | 5i/L/A (N2B27-based) | PD0325901, IM-12, SB590885, WH-4-023, Y-27632, LIF, Activin A | hESC.h9_n1 |
| | | WIBR3 | 5i/L/A (N2B27-based) | | hESC.38wt_n1 |
| | | H9 | a5i/L/A (N2B27-based) | AZ628, IM-12, SB590885, WH-4-023, Y-27632, LIF, Activin A | hESC.h9_n3 |
| | | WIBR3 | a5i/L/A (N2B27-based) | | hESC.38wt_n3 |
| | (+)PKCi | H9 | AXGY | AZ628, XAV939, Go6983 (PKC inhibitor), Y-27632 | hESC.h9_n6 |
| | | WIBR3 | AXGY | | hESC.38wt_n6 |
| | | H9 | AXGYU | AZ628, XAV939, Go6983 (PKC inhibitor), Y-27632, UNC0638 | hESC.h9_n7 |
| | | WIBR3 | AXGYU | | hESC.38wt_n7 |

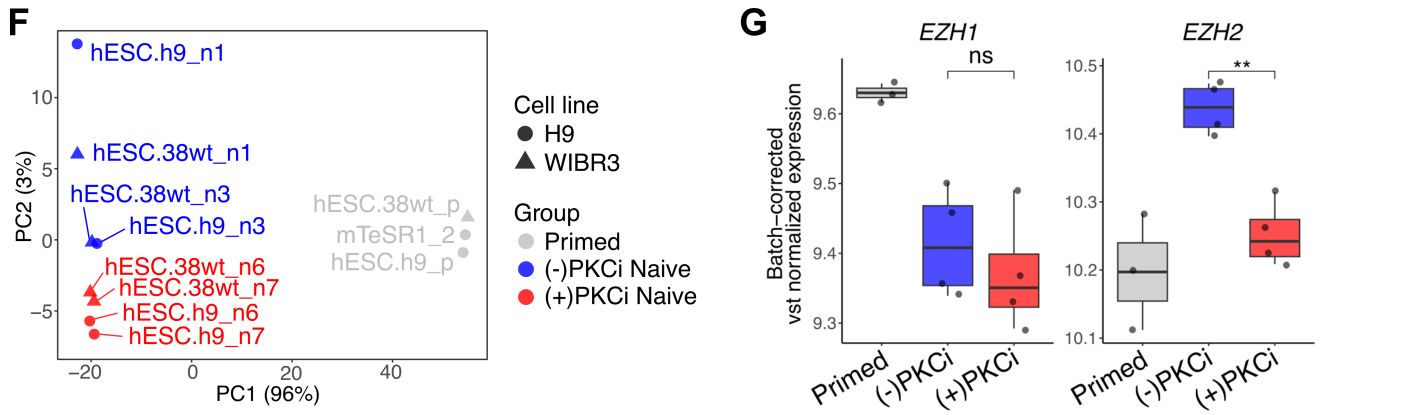
